# Chromatin interactome mapping at 139 independent breast cancer risk signals

**DOI:** 10.1101/520916

**Authors:** Jonathan Beesley, Haran Sivakumaran, Mahdi Moradi Marjaneh, Luize G. Lima, Kristine M. Hillman, Susanne Kaufmann, Natasha Tuano, Nehal Hussein, Sunyoung Ham, Pamela Mukhopadhyay, Stephen Kazakoff, Jason S. Lee, Kyriaki Michailidou, Daniel R. Barnes, Antonis C. Antonio, Laura Fachal, Alison M. Dunning, Douglas F. Easton, Nicola Waddell, Joseph Rosenbluh, Andreas Möller, Georgia Chenevix-Trench, Juliet D. French, Stacey L. Edwards

**Affiliations:** Cancer Program, QIMR Berghofer Medical Research Institute, Brisbane, Australia; Department of Biochemistry and Molecular Biology, Monash University, Melbourne, Australia; Faculty of Medicine, The University of Queensland, Brisbane, Australia; Centre for Cancer Genetic Epidemiology, Department of Public Health and Primary Care, University of Cambridge, Cambridge, UK; Department of Electron Microscopy/Molecular Pathology, The Cyprus Institute of Neurology and Genetics, Nicosia, Cyprus; Centre for Cancer Genetic Epidemiology, Department of Oncology, University of Cambridge, Cambridge, UK

**Author notes:** These authors contributed equally (Jonathan Beesley, Haran Sivakumaran, Mahdi Moradi Marjaneh). Senior author (Juliet D. French, Stacey L. Edwards). Correspondence (S.L.E) (J.D.F).

## Abstract

Genome-wide association studies have identified 196 high confidence independent signals associated with breast cancer susceptibility. Variants within these signals frequently fall in distal regulatory DNA elements that control gene expression. We designed a Capture Hi-C array to enrich for chromatin interactions between the credible causal variants and target genes in six human mammary epithelial and breast cancer cell lines. We show that interacting regions are enriched for open chromatin, histone marks for active enhancers and transcription factors relevant to breast biology. We exploit this comprehensive resource to identify candidate target genes at 139 independent breast cancer risk signals, and explore the functional mechanism underlying altered risk at the 12q24 risk region. Our results demonstrate the power of combining genetics, computational genomics and molecular studies to rationalize the identification of key variants and candidate target genes at breast cancer GWAS signals.

## INTRODUCTION

Breast cancer is known to have an important inherited component. While rare coding mutations in susceptibility genes such as *BRCA1, BRCA2* and *PALB2* confer a high risk of breast cancer, these account for less than one quarter of the familial risk^1^. Much of the remaining heritability is due to the combination of a large number of common, low-penetrance variants^2,3^. Genome-wide association studies (GWAS) have been a powerful tool to identify disease-associated genetic variants, but these studies do not directly address the underlying biological mechanisms. A combination of fine scale-mapping, bioinformatic and functional studies are required to establish this link^4^. The Breast Cancer Association Consortium (BCAC) and the Consortium of Investigators of Modifiers of *BRCA1/2* (CIMBA) have recently performed large-scale genetic fine-mapping of 150 breast cancer susceptibility regions in ∼217,000 breast cancer cases and controls of European ancestry^5^. Step-wise multinomial logistic regression analysis identified 196 high confidence independent risk signals, defined as having association p values < 10^-6^ after adjusting for other variants. Fachal et al (2018) used these data to define sets of credible causal variants (CCVs) for each signal, defined as variants with p values within two orders of magnitude of the top variant.

The majority of CCVs mapped to non-protein-coding regions of the genome and are enriched at regulatory DNA elements such as enhancers, silencers and insulators^2,5^. It is established that many regulatory elements are located long distances from their target gene promoters, and that regulation of transcription involves direct physical interactions brought about by chromatin looping^6^. Importantly, individual enhancers often loop to and regulate multiple genes, including protein-coding and noncoding RNA genes. Adding to the complexity, enhancers do not necessarily act on the closest promoter but can bypass neighbouring genes to regulate genes located more distally. There is also considerable evidence that most enhancer-promoter interactions occur in *cis* and within chromatin structures called topologically associating domains (TADs)^7^. TADs are typically several hundred kilobases to a few megabases in size and are relatively stable between cell types and in response to extracellular signals^8,9^.

Various chromatin conformation capture (3C)-based methods have been developed to map chromatin contacts at a genome-wide level. The basic principle of 3C involves chromatin fragmentation of formaldehyde-fixed nuclei (usually by restriction digestion), followed by ligation of linked DNA fragments, then detection and quantification of ligation products^10^. One of these methods, Hi-C, is an unbiased but relatively low-resolution approach, that quantifies interactions between all possible DNA fragment pairs in the genome^11^. Hi-C has been used extensively to analyze the three-dimensional organization of genomes, including compartmentalization of chromatin and the position of TADs^12,13^. To increase Hi-C resolution, several groups have developed sequence capture to enrich for chromosomal interactions involving targeted regions of interest^14^-^17^. There are several capture methodologies, but typically RNA or DNA oligonucleotide baits are directed to the ends of targeted DNA fragments to enrich for ligation events prior to next generation sequencing^18,19^. Promoter Capture Hi-C (PCHi-C) is the most widely used approach where baits are designed to annotated promoters, resulting in strong enrichment for promoter-anchored interactions^15^-^17,20^. A few post-GWAS studies have also used Region Capture Hi-C, in which baits target linkage disequilibrium blocks or restriction fragments containing genetic variants associated with the disease of interest^21,22^.

Here, we applied Variant Capture Hi-C (VCHi-C) and PCHi-C to normal breast and breast cancer cell lines to generate a catalog of interactomes. We report several hundred candidate target genes in breast cancer risk regions including some known cancer driver genes but also many molecular targets not previously implicated in breast cancer etiology.

## RESULTS

### VCHi-C and PCHi-C interaction profiling

To enrich for chromatin interactions relevant to breast cancer risk, we designed two capture arrays, Variant Capture (VC) and Promoter Capture (PC). The VCHi-C baits were designed to *HindIII* fragments that contained at least one CCV, regardless of the CCV regulatory potential (**Figure 1A**;^5^). We could design baits to 190/196 signals (97%) which included 6044/7394 CCVs. The PCHi-C baits were designed to annotated promoters within 1 Mb of CCVs at breast cancer risk signals (**Figure 1A**). This dual-capture approach ensured comprehensive coverage of each risk signal and provided independent validation of interactions. We performed *in situ* VCHi-C and PCHi-C^16,18^ in two non-tumorigenic breast cell lines (B80T5, MCF10A), two estrogen receptor positive (ER+; MCF7, T47D) and two ER-(MDAMB231, Hs578T) breast cancer cell lines. Sequencing of both captures produced over one billion unique di-tags involving CCV-containing fragments and annotated promoters (**Table S1**). To assess the robustness of the approach, each CHi-C experiment was conducted in two biological replicates per cell type. We observed strong correlation between the replicates, particularly when captured interaction pairs were within 0.5 Mb (**Figure S1A**).

**Figure 1.**
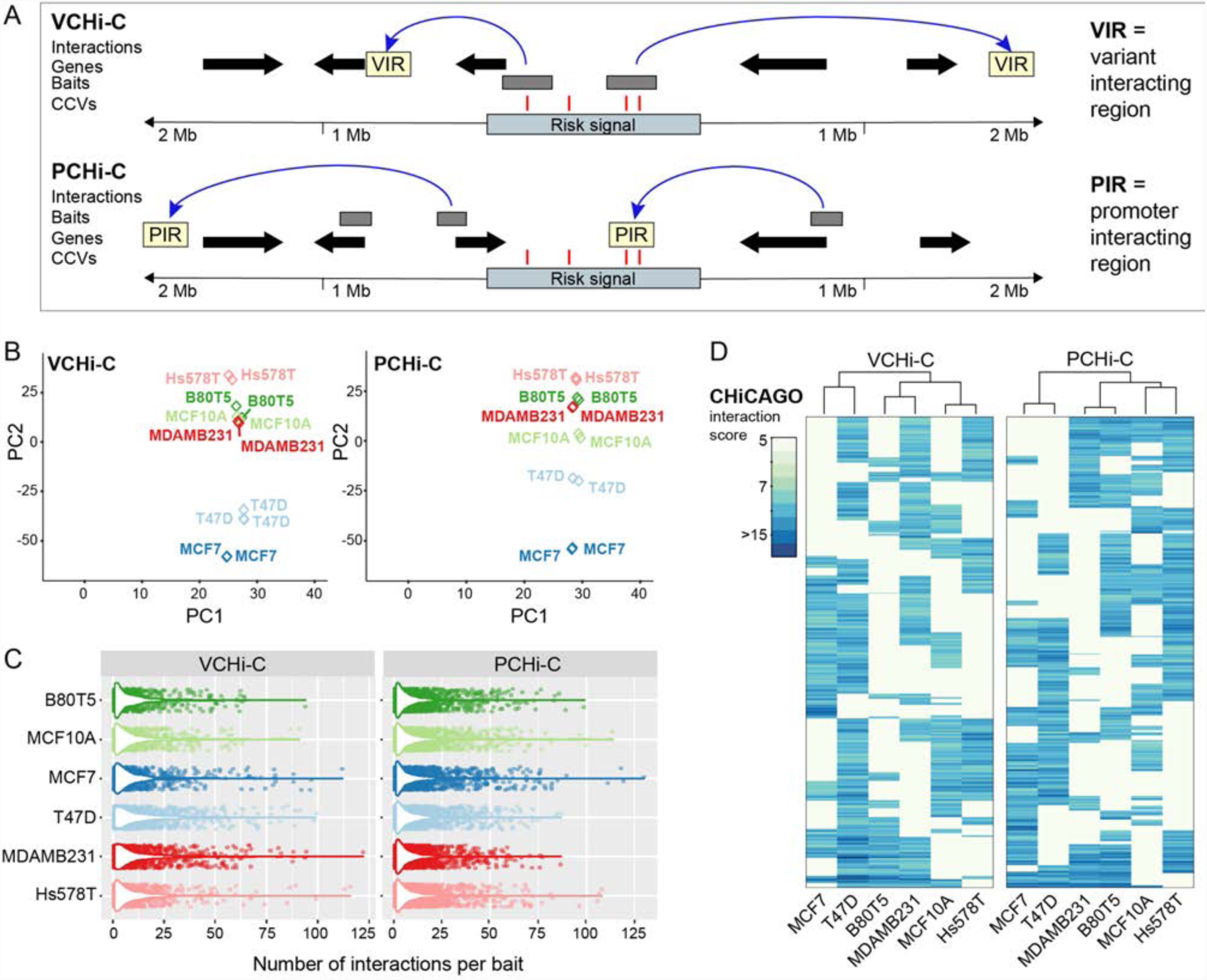
VCHi-C and PCHi-C in human breast cell lines. **(a)** Schematic of a hypothetical breast cancer risk signal and plausible chromatin interactions. Chromatin interactions are shown as blue arcs. Genes are depicted as black arrows. CHi-C baits are depicted as gray boxes. CCVs are shown as red vertical lines. The colored boxes illustrate variant-interacting regions (VIRs) or promoter-interacting regions (PIRs). **(b)** Principle component analysis of CHiCAGO-scored interactions in VCHi-C or PCHi-C biological replicates. **(c)** Distribution of CHiCAGO-scored interaction number per bait per cell line (combined biological replicates). **(d)** Agglomerative hierarchical clustering for the VCHi-C and PCHi-C in six breast cell lines.

We initially used the CHiCAGO pipeline^23^ to assign confidence scores to interactions derived from the VCHi-C and PCHi-C (**Table S2**). Principal component analysis (PCA) based on CHiCAGO scores demonstrated concordance for individual replicates in the VCHi-C and PCHi-C. PCA was able to separate ER+ breast cancer from normal breast or ER-breast cancer cell lines (**Figure 1B**). Using a strict interaction threshold (CHiCAGO score ≥?5, intrachromosomal and interaction distance ≤2Mb) we detected on average ∼10,000 VCHi-C and ∼27,000 PCHi-C high-confidence interactions per cell type (**Figure 1C** and **Table S2**). The difference in interaction number between captures likely reflects the higher number of PCHi-C baits. In addition, VCHi-C baits were designed to all possible CCV-containing *HindIII* fragments, but some CCVs will be correlated passenger variants or function through alternative non-looping mechanisms, such as promoter variants. For the VCHi-C, we detected a median of five variant-interacting regions (VIRs; **Figure 1A**) per bait per cell type, of which 3-5% interacted with an annotated protein-or non-coding promoter. Similarly, for the PCHi-C, we detected a median of five promoter-interacting regions (PIRs; **Figure 1A**) per bait per cell type, where 2.4% specifically interacted with a CCV-containing fragment (**Figure S1B** and **Table S2**). The median linear distance between interactions from either capture ranged from 192-405 kb (**Figure S1C**) and ∼70% of the CHi-C interactions occurred within TAD boundaries. Hierarchical clustering based on CHiCAGO scores separated the cells lines based on ER-status (**Figure 1D**), which suggested that ER status mediates cell-type specificity of the interactomes. We also observed a positive correlation (Pearson’s r = 0.60-0.84) in CHiCAGO scores for interactions detected in both the VCHi-C and PCHi-C (**Figure S1D**), thus validating our approach.

### Interacting regions are enriched for regulatory features, eQTLs and CCVs in breast cells

We first annotated CHiCAGO-scored PIRs in each breast cell type with DNase-seq data derived from a diverse panel of cells and tissues as part of the Roadmap Epigenomics Project^24^. We found PIRs to be enriched for regions of accessible chromatin in human mammary epithelial cells (HMEC), as compared to non-breast cells (**Figures 2A** and **S2A**). To explore this observation in additional breast cells, we annotated PIRs with assay for transposase-accessible chromatin sequencing (ATAC-seq) peaks in five breast cell lines (**Table S3**) and noted that the enrichment signals were strongest from PIRs detected in the matched cell line (**Figure 2B**). We next investigated the epigenetic makeup of PIRs using ChIP-seq data for histone modifications and other DNA-binding proteins in human cell lines. PIRs were significantly enriched for histone marks associated with active enhancers (H3K27ac and H3K4me1) as compared to inactive elements which are typically marked by the polycomb-associated mark H3K27me3 (**Figure 2C**).

**Figure 2.**
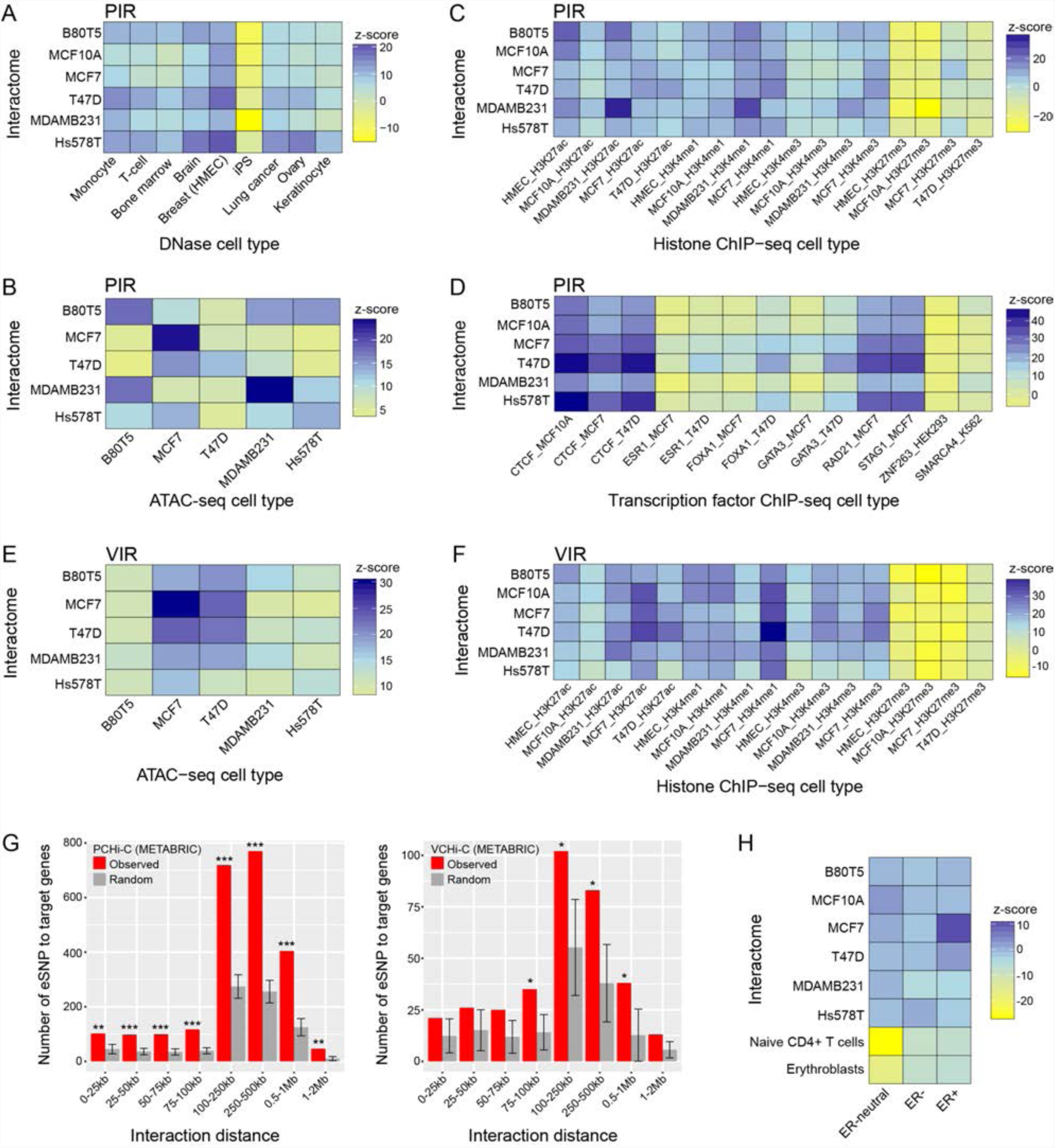
PIRs and VIRs share regulatory features. Heatmaps showing promoter-interacting region (PIR) enrichment for **(a)** DNase I hypersensitivity sites in a diverse range of cell types, **(b)** ATAC-seq peaks in breast cell lines, **(c)** histone marks by ChIP-seq in available breast cell lines, and **(d)** relevant transcription factor binding in available breast cell lines, expressed as z-scores. Heatmaps showing variant-interacting region (VIR) enrichment for **(e)** ATAC-seq peaks in breast cell lines and **(f)** histone marks by ChIP-seq in available breast cell lines, expressed as z-scores. **(g)** The number of interactions between expression single nucleotide polymorphisms (eSNPs) and associated target genes (observed) compared to randomly assigned interactions (random), binned by interaction distance. Asterisks represent the significance of enrichment of observed versus randomized interacting regions (permutation test *p<0.05, **p<0.01, ***p<0.001). **(h)** Heatmap showing CCV enrichment in PIRs in breast cell lines. CCVs are classified as conferring greater risk of developing ER+, ER-or both (ER-neutral) tumor subtypes.

Binding sites for several structural proteins with established roles in chromatin looping were also enriched in PIRs, including CTCF and the cohesin subunits RAD21 and STAG1 (**Figure 2D**), consistent with the role of these factors in mediating long-range genomic interactions^9,12^. Associations were also observed for the cistromes of important breast cancer transcription factors (TFs); ESR1, FOXA1 and GATA3 (**Figure 2D**). This enrichment was stronger in the ER+ MCF7 (z-score=5.04) and T47D (z-score=2.97) cell lines as compared to available ER-breast cancer, normal breast and other non-breast cell lines (**Figure S2B**), consistent with an additional layer of ER-mediated cell-type specificity^25^. Applying the same enrichment criteria, we also found VIRs to be enriched in ATAC-seq peaks in the matched cell associated with active enhancers (H3K27ac and H3K4me1) and also for H3K4me3, which marks active gene promoters (**Figure 2F**), supporting the notion that promoters and enhancers cooperatively communicate through transcriptionally active chromatin^26^.

To demonstrate PIR and VIR gene regulatory function, we assessed the overlap of expression quantitative trait loci (eQTLs) in normal breast tissue from the METABRIC (Molecular Taxonomy of Breast Cancer International Consortium) cohort^2,27^. We found 800 eQTL genes (eGenes) with eSNPs (false discovery rate (FDR)<0.05) within PIRs in at least one analyzed breast cell line. Examination of the VIR data also revealed 184 eGenes interacting with eSNPs (**Figure 2G**). To assess specificity of eQTL localization to interacting regions, we maintained the interaction network by assigning baits to randomly selected promoters and compared the number of interactions supported by eQTL-target gene pairs. We found that eQTLs were significantly more likely to loop to their associated gene than expected by chance, across a broad range of linear distances from their target promoters (**Figure 2G**). Finally, we integrated the PIRs with CCVs^5^ and found that CCVs stratified for their association with ER+ and/or ER-tumor subtypes were enriched at PIRs in the ER+ breast cancer cell lines (**Figure 2H**). This enrichment was not as pronounced for the ER-and ER-neutral CCVs in the ER-breast cancer and normal breast cell lines, which may indicate a lack of statistical power to detect enrichment or that the underlying mechanisms are more heterogeneous.

### Fine-mapping of VCHi-C and PCHi-C profiles

While the CHiCAGO pipeline is extremely useful for interaction detection in CHi-C data^23^, many of the generated contact maps contain contiguous restriction fragments linked with the same target. It is hypothesized that such collateral contacts might result from inaccuracy during the cross-linking process in CHi-C^28^ or from bait migration via Brownian motion^29^. Therefore, as a complementary interaction scoring method, we also used a recently developed Bayesian sparse variable selection approach (“*Peaky*”; ^30^). The model proposes that for any given bait, the expected CHi-C signal at each prey fragment is expressed as a sum of contributions from a set of fragments directly contacting that bait^30^. We applied Peaky to the ∼1300 baits from the VCHi-C and ∼3200 baits from the PCHi-C (**Table S4**) to derive a measure of confidence in the location of a direct contact called the marginal posterior probability of a contact (MPPC)^30^.

To facilitate a comparison with CHiCAGO-scored interactions, we applied an interaction threshold of MPPC ≥0.1. We filtered for intrachromosomal and interaction distance ≤2 Mb and detected ∼3,500 VCHi-C and ∼7,400 PCHi-C interactions per cell type (**Figure S3A** and **Table S4**). For the VCHi-C, ∼11% of CCV-containing fragments interacted with an annotated protein-or non-coding promoter and for the PCHi-C, ∼2.5% of promoter fragments specifically interacted with a CCV-containing fragment (**Figure S3B** and **Table S4**). There were fewer interactions detected by Peaky, perhaps because Peaky can distinguish and rank a subset of direct contacts from long stretches of chromatin interactions^30^. The median linear distance between interactions from either capture was longer than CHiCAGO-scored interactions (ranged from 294-489 kb; **Figure S3C**). Similar to CHiCAGO-scored interactions, hierarchical clustering based on MPPC scores also separated the cell lines based on ER status (**Figure S3D**). We then compared the CHiCAGO and MPPC scores for each bait-prey pair. As reported by Eijsbouts et al^30^, we noted that CHiCAGO and MPPC scores were positively correlated (**Figure S3E**; Spearman’s ?ρ = 0.22-0.37). Peaky was able to refine the number of CHiCAGO-scored interactions by 12-17% in both captures; however a proportion of interactions were identified by Peaky but not CHiCAGO (**Figure S3F**). To provide a more stringent list of CCVs and candidate target genes, we combined inferences from the two approaches.

### Prioritizing CCVs by Peaky fine-mapping of the PCHi-C data

At many signals, we noted that CHiCAGO identified long stretches of PIRs, some of which contained CCVs. We therefore used Peaky to fine-map the CHiCAGO identified interactions to identify the likely driver contacts within these stretches. This approach proved particularly useful at 9q33.1, where CHiCAGO identified 24 PIRs starting at ∼340 kb from a *PAPPA* (Pregnancy-associated plasma protein A) promoter (**Figure 3A**). Peaky fine-mapping using a *PAPPA* promoter bait indicated this stretch of interactions might be explained by a subset of contacts (MPPC ≥ 0.1), which spanned one (rs811688) out of 29 CCVs in MCF7 cells (**Figure 3A**). 3C provided further support that the *HindIII* fragment containing rs811688 was the most frequently interacting fragment with the *PAPPA* promoter (**Figure S4A**). *PAPPA* encodes a secreted zinc metalloproteinase and is an important regulatory component of the insulin-like growth factor system^31^. Recent studies indicate PAPPA is frequently overexpressed in luminal B breast tumors^32^ and identify PAPPA as a pregnancy-dependent oncogene that promotes the formation of pregnancy-associated breast cancer^33^.

**Figure 3.**
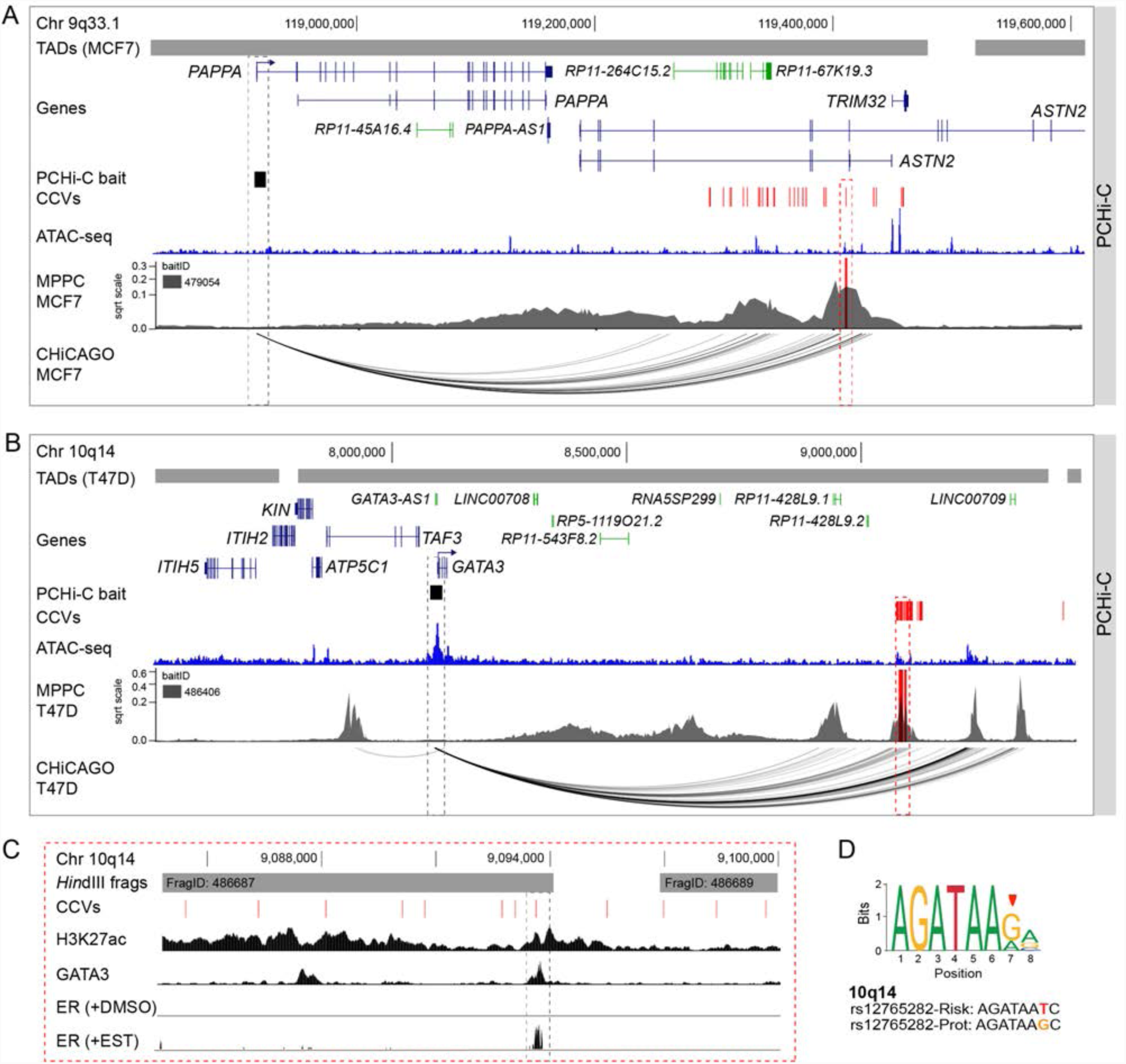
PCHi-C Peaky fine-mapping prioritizes CCVs at 10q14 and 6p22.3. **(a)** Chromatin interactions at 9q33.1 in MCF7 breast cancer cells. Topologically associating domains (TADs; **Table S7**) are shown as horizontal gray bars above GENCODE annotated coding (blue) and non-coding (green) genes. The PCHi-C bait is depicted as a black box. CCVs are shown as red vertical lines. The ATAC-seq track is shown as a dark blue histogram. Peaky defined MPPC values (from PCHi-C BaitID: 479054) are plotted with the prioritized CCV overlaid as a red vertical line. CHiCAGO-scored interactions are shown as black arcs. The dashed red outline highlights the prioritized CCV rs811688 and the dashed gray outline the target gene (*PAPPA*). **(b)** Chromatin interactions at 10q14 in T47D breast cancer cells. Topologically associating domains (TADs) are shown as horizontal gray bars above GENCODE annotated coding (blue) and non-coding (green) genes. The PCHi-C bait is depicted as a black box. CCVs are shown as red vertical lines. The ATAC-seq track is shown as a dark blue histogram. Peaky defined MPPC values (from PCHi-C BaitID: 486406) are plotted with the prioritized CCVs overlaid as red vertical lines. CHiCAGO-scored interactions are shown as black arcs. The dashed red outline highlights the prioritized CCVs and the dashed gray outline the target gene (*GATA3*). **(c)** Zoomed in view of prioritized CCVs at 10q14. *HindIII* fragments are shown as gray bars with their fragment IDs. CCVs are shown as red vertical lines. Black histograms denote ChIP-seq data from T47D cells for H3K27ac, GATA3 and estrogen receptor (ER; cells treated with DMSO or 17 beta-estradiol (EST)). The dashed gray outline highlights CCV rs12765282. **(d)** Position weight matrix of the GATA3 binding site from JASPAR (red arrowhead indicates the CCV position in the motif), with homology to the risk (*t*) and protective (*g*) alleles of rs12765282 colored below.

Another example is 10q14, where CHiCAGO identified 59 PIRs located ∼1 Mb from the *GATA3* (GATA binding protein 3) promoter. Interactions between *GATA3* and CCVs were restricted to the ER+ (T47D and MCF7) breast cell lines and spanned 49 CCVs (**Figure 3B**). Peaky fine-mapping indicated this stretch of interactions might be explained by a subset of four contacts, one of which spanned a region containing CCVs. Two *HindIII* fragments within the CCV-containing peak surpassed the 0.1 MPPC interaction threshold and contained 11 out of the 49 CCVs (**Figure 3B**). 3C provided further support that the *HindIII* fragment containing 8 CCVs (FragID: 486687) was the most frequently interacting fragment with the *GATA3* promoter (**Figure S4B**). Notably, one CCV (rs12765282) within the 3C-identified peak mapped to a putative regulatory element as defined by H3K27ac marks and TF binding in T47D cells (**Figure 3C**). This CCV is predicted to alter a GATA3-binding motif, with the risk allele likely acting to decrease GATA3 binding. ChIPseq data showed that GATA3 and ER bound to the CCV site in T47D cells, which are homozygous for the protective *g*-allele (**Figures 3C** and **3D**). GATA3 is important in mediating enhancer accessibility for ER^25^, raising the possibility of a GATA3-mediated regulatory loop underlying risk at this region.

Taken together, at 77 signals where we could detect at least one promoter-CCV interaction, we could prioritize 839 out of 4208 CCVs using the combined CHICAGO (score ≥?5) and Peaky (MPPC ? 0.1) fine-mapping approach. This included 33 signals where the number of prioritized genetically indistinguishable CCVs could potentially be reduced to less than five at each signal (**Table S5**).

### Prioritizing target genes by sequential CHiCAGO and Peaky fine-mapping

The combined analyses can be extended to integrate, where possible, the VCHi-C and PCHi-C data. One example is 1p22.3, where CHiCAGO detected interactions in the VCHi-C data between two independent signals and the *LMO4* (LIM-only protein 4) promoter in Hs578T breast cancer cells (**Figure 4A**). Peaky fine-mapping using signal 2 VCHi-C baits then provided further support that *LMO4* was the likely target gene (**Figure 4A**). Peaky was also applied to signal 1 VCHi-C baits, but the resulting contact peaks did not reach the 0.1 MPPC interaction threshold (**Figure S4C**). We then interrogated the PCHi-C data using two *LMO4* promoter baits in Hs578T cells. CHiCAGO identified 84 PIRs starting at ∼612 kb from the *LMO4* promoter (**Figure 4A**). Peaky fine-mapping using the same promoter baits indicated this stretch of interactions might be explained by a subset of three direct contacts (MPPC ?0.1). One contact spanned two *HindIII* fragments within signal 2 and potentially prioritized four out of eight CCVs at this signal (**Figure 4B**). Of these, one CCV (rs3008455) mapped to a putative regulatory element as defined by open chromatin and TF binding in normal breast cells (**Figure 4B**). This CCV is predicted to alter a CTCF-binding motif, with the risk allele promoting increased CTCF binding (**Figure 4C**). LMO4 is a transcriptional modulator that is overexpressed in >50% of breast tumors^34^. Overexpression of LMO4 promotes cell proliferation, invasion and tumor formation and induces mammary hyperplasia in transgenic mice^35^.

**Figure 4.**
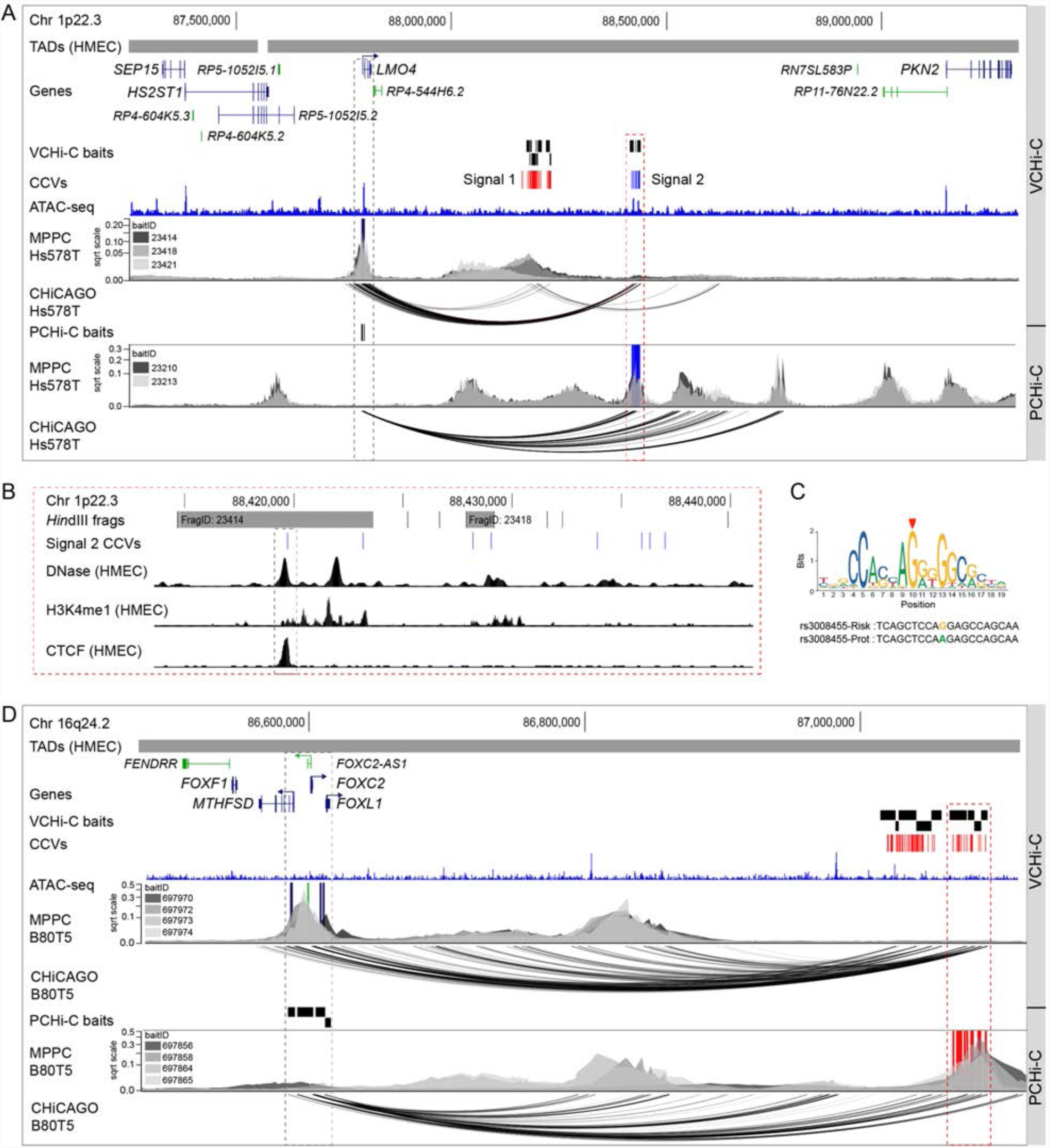
Sequential CHiCAGO and Peaky fine-mapping prioritizes CCVs and target genes. **(a)** Chromatin interactions at 1p22.3 in Hs578T breast cancer cells. Topologically associating domains (TADs) are shown as horizontal gray bars above GENCODE annotated coding (blue) and non-coding (green) genes. The VCHi-C or PCHi-C baits are depicted as black boxes. Risk signals 1 and 2 are numbered and the CCVs within each signal are shown as colored vertical lines. The ATAC-seq track is shown as a dark blue histogram. Peaky defined MPPC values (from specified BaitIDs) are plotted with the prioritized gene overlaid as a dark blue vertical line or prioritized CCVs overlaid as royal blue vertical lines. CHiCAGO-scored interactions for specified BaitIDs are shown as black arcs. The dashed red outline highlights the prioritized CCVs and the dashed gray outline the target gene (*LMO4*). **(b)** Zoomed in view of prioritized signal 2 CCVs at 1p22.3. VCHi-C baits are shown as gray bars with their fragment IDs. CCVs are shown as blue vertical lines. Black histograms denote DNase I hypersensitivity sites or ChIP-seq data for H3K4me1 and CTCF binding from HMEC cells. The dashed gray outline highlights CCV rs3008455. **(c)** Position weight matrix of the CTCF binding site from JASPAR (red arrowhead indicates the CCV position in the motif), with homology to the risk (*g*) and protective (*a*) alleles of rs3008455 colored below. **(d)** Chromatin interactions at 16q24.2 in B80T5 normal breast cells. Topologically associating domains (TADs) are shown as horizontal gray bars above GENCODE annotated coding (blue) and non-coding (green) genes. The VCHi-C or PCHi-C baits are depicted as black boxes. CCVs are shown as red vertical lines. The ATAC-seq track is shown as a dark blue histogram. Peaky defined MPPC values (from specified BaitIDs) are plotted with the prioritized genes overlaid as dark blue or green vertical lines and prioritized CCVs overlaid as red vertical lines. CHiCAGO-scored interactions for specified BaitIDs are shown as black arcs. The dashed red outline highlights the prioritized CCVs and the dashed gray outline the prioritized target genes (*MTHFSD, FOXC2, FOXL1* and *FOXC2-AS1*).

A more complex example is 16q24.2, where CHiCAGO detected 62 VIRs spanning a ∼320 Kb genomic region derived from nine separate VCHi-C baits (**Figure 4D**). Peaky fine-mapping of this VCHi-C data then prioritized *FOXC2, FOXC2-AS1, FOXL1* and *MTHFSD* as the likely target genes in B80T5 normal breast cells (**Figure 4D**). We interrogated the PCHi-C data using the four target gene promoter baits in B80T5 cells. CHiCAGO identified 40 PIRs spanning a ∼500 Kb genomic region. Peaky fine-mapping using the same promoter baits indicated this stretch of interactions might be explained by a subset of two direct contacts (MPPC ≥0.1). One contact spanned five *HindIII* fragments and potentially prioritized 21 out of the possible 85 CCVs at this signal (**Figure 4D**). Preliminary *in silico* analyses revealed many of the 21 prioritized CCVs display regulatory activity and therefore additional studies would be required to determine which are the likely functional variants. FOXC2 and FOXL1 are members of the Forkhead family of transcription factors with important functions in biological processes such as cell cycle control, proliferation and development^36^. FOXC2 has been implicated in triple-negative breast cancer progression and therapy resistance^37^, while FOXL1 is reported to inhibit breast cancer cell proliferation, invasion, and migration^38^. Little is known about *MTHFSD* (Methenyltetrahydrofolate synthetase domain-containing), but a recent report suggests the gene encodes a stress granule-associated RNA-binding protein^39^.

### Identification of 651 candidate target genes at 139 breast cancer risk signals

We defined candidate target genes of breast cancer risk signals by CHiCAGO-and/or Peaky-scored CCV-gene promoter interactions in VCHi-C or PCHi-C in at least two cell lines. This combined analysis resulted in 651 candidate target genes at 139 breast cancer risk signals, including 419 protein-coding genes (**Table S5**). The majority of candidate target genes interacted with one signal, but ∼13% interacted with two or more independent signals (**Figure S4D**). The 6q25 region is one of the more extreme examples, where five out of six independent signals all loop to and potentially regulate *ESR1* (**Figure S4E).** More than 80% of signal-target gene interactions skipped at least one annotated gene promoter and ∼75% of signals interacted with at least two promoter-containing fragments (**Figures S4F**). One example that demonstrates both characteristics is 8q24.13, where signal 1 CCVs interact with six candidate target genes (*WDYHV1, FBXO32, CTD-2552K11.2, ANXA13, FAM91A1* and *TRMT12*) including skipping three annotated genes to contact the *TRMT12* promoter (**Figure S4G**). Notably, 181 candidate target genes were identified by both CHiCAGO and Peaky (**Figure S4H**), which may further prioritize these genes for functional validation. This priority list includes established breast cancer driver genes such as *MYC* and *GATA3*^40^, but also includes many genes with no reported role in breast cancer (**Table S5**).

### CHi-C identifies *TBX3* as the target of multiple risk signals

To further illustrate the power of combining genetic fine-mapping, CHi-C and functional studies, we examined in detail the 12q24 susceptibility region. Genetic fine-mapping of 12q24 identified at least four independent signals^2,5^ (listed in **Table S6**); signal 1 (seven CCVs), signal 2 (one CCV) and signal 4 (six CCVs) were more strongly associated with ER+ tumors, whereas signal 3 (eight CCVs) was associated with both ER+ and ER-breast cancer (**Table S6**). The CCVs in all four signals are located in a large intergenic region on 12q24 between *TBX3* and *MED13L* (**Figure 5A**). We used ATAC-seq and available ChIP-seq datasets from ENCODE^41^ to map CCVs relative to transcriptional regulatory elements. This analyses showed evidence of putative regulatory elements overlapping the CCVs at each signal, indicating that one or more CCVs likely have high regulatory potential (**Figure 5A**). CHi-C and 3C identified *TBX3* (T-Box 3) as the most likely target gene (**Figures 5A, S5A** and **Table S6**). Notably, we detected interactions between *TBX3* and each of the four independent signals in a cell-type specific manner (**Figures 5A** and **S5B**).

**Figure 5.**
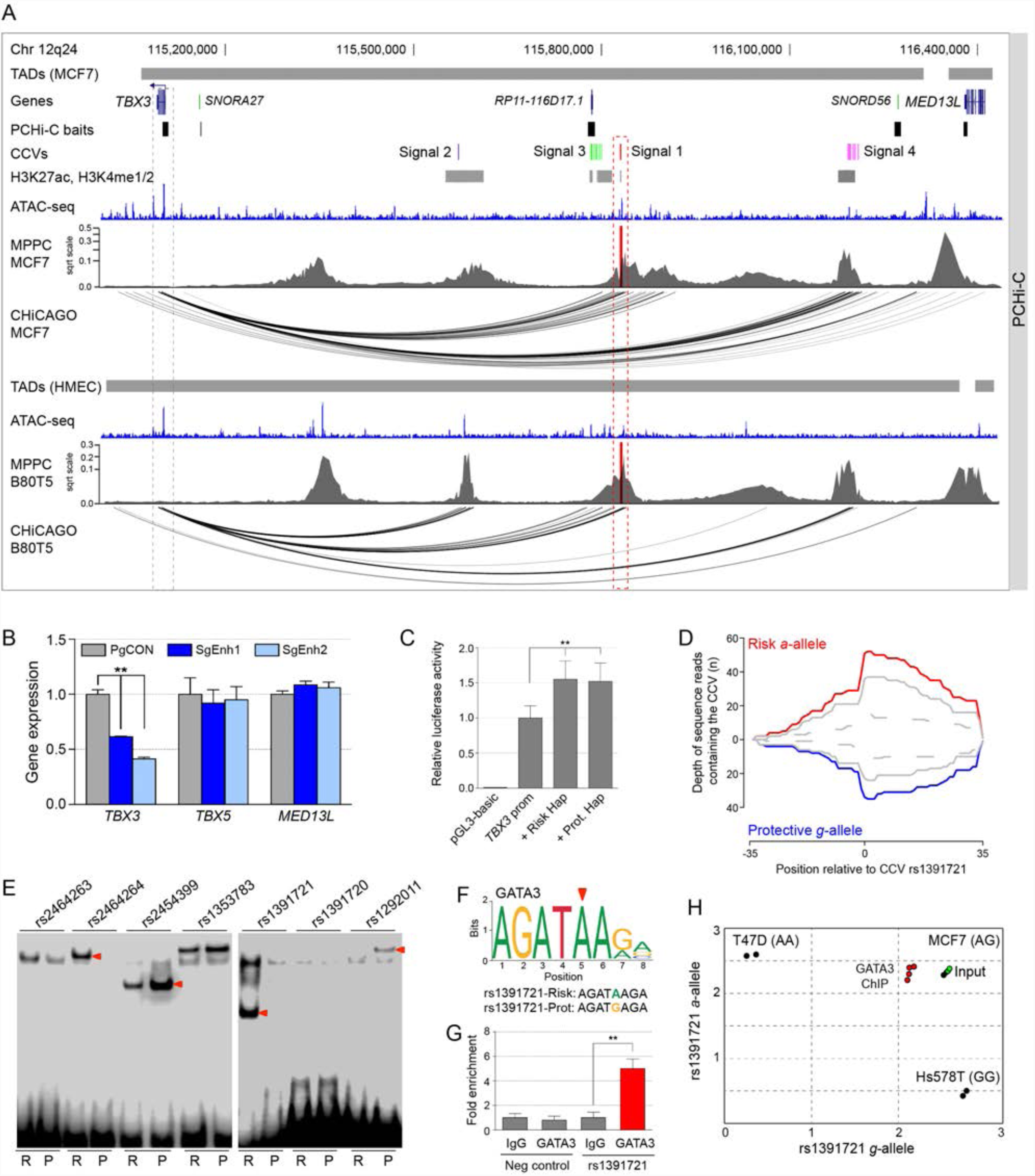
Molecular analysis of signal 1 CCVs at 12q24. **(a)** Chromatin interactions in MCF7 and B80T5 breast cell lines. Topologically associating domains (TADs) are shown as horizontal gray bars above GENCODE annotated coding (blue) and non-coding (green) genes. The PCHi-C baits are depicted as black boxes. Risk signals 1-4 are numbered and the CCVs within each signal are shown as colored vertical lines. ENCODE ChIP-seq data for available histone marks are depicted as gray boxes. The ATAC-seq tracks are shown as dark blue histograms. Peaky defined MPPC values (from PCHi-C BaitID: 596031) are plotted with the prioritized CCVs overlaid as red vertical lines. CHiCAGO-scored interactions are shown as black arcs. The dashed red outline highlights signal 1 CCVs and the dashed gray outline the target gene (*TBX3*). **(b)** The 12q24 enhancer was repressed by targeting dCas9-KRAB to the enhancer in MCF7 cells with two different CRISPRi single-guide (sg) RNAs (SgEnh1 and SgEnh2). PgCON contains a non-targeting control sgRNA. Gene expression of *TBX3, TBX5* and *MED13L* was measured by qPCR and normalized to *GUSB*. Error bars represent the SEM (n = 3). p values were determined by two-way ANOVA followed by Dunnett’s multiple-comparison test (**p<0.01). **(c)** Luciferase reporter assays following transient transfection of MCF7 cells. The 12q24 enhancer containing either the risk or protective (Prot.) haplotype was cloned into *TBX3* promoter-driven luciferase constructs (*TBX3* prom). Error bars represent the SEM (n = 3). p values were determined by two-way ANOVA followed by Dunnett’s multiple-comparison test (**p<0.01). **(d)** Allele-specific DNase I hypersensitivity at CCV rs1391721 in heterozygous MCF7 cells. The depth of reads containing the risk (red) and protective (blue) alleles are shown. **(e)** EMSAs for signal 1 CCVs to detect allele-specific binding of nuclear proteins. Labeled oligonucleotide duplexes were incubated with MCF7 nuclear extract. Red arrowheads show bands of different mobility detected between risk (R) and protective (P) alleles. **(f)** Position weight matrix of the GATA3 binding site from JASPAR, with homology to the risk (*a*) and protective (*g*) alleles of rs1391721 colored below. **(g)** Allele-specific GATA3 ChIP-PCR results assessed at CCV rs1391721 in heterozygous MCF7 cells. Error bars represent the SEM (n = 3). p values were determined by a two-tailed Student’s *t*-test (**p<0.01). **(h)** Allelic discrimination plot of the GATA3 ChIP in MCF7 cells. Genomic DNA extracted from homozygous T47D and Hs578T breast cancer cells were used as controls.

Our functional studies focused on the strongest signal 1 CCVs. CRISPRi-silencing of the signal 1 element in ER+ MCF7 cells showed that *TBX3*, but not *TBX5* and *MED13L*, levels were significantly reduced (**Figure 5B**). Reporter assays then confirmed that the element acts as an enhancer on the *TBX3* promoter in the presence of either the risk or protective haplotypes (**Figure 5C**). We used available DNase-seq data derived from heterozygous MCF7 cells to show that the risk *a*-allele of CCV rs1391721 may promote allele-specific open chromatin (**Figure 5D**). Electrophoretic mobility shift assays (EMSAs) then assessed TF binding for each of the signal 1 CCVs. Allele-specific binding by nuclear proteins was observed for CCVs rs2464264, rs2454399, rs1391721 and rs1292011 in MCF7 and BT474 extracts (**Figures 5E** and **S6A**). Further EMSAs using competitor DNA against predicted TFs suggested GATA3 bound to the rs1391721 site (**Figure S6B**). Similar to the 10q14 CCV, rs1391721 is also predicted to lie in a GATA3 binding site. Here, the risk *a*-allele promoted increased GATA3-binding compared to the protective *g*-allele (**Figure 5F**), as evident in GATA3 ChIP-seq data derived from heterozygous MCF7 cells (**Figure S6C**). To assess occupancy of GATA3 *in vivo*, we performed ChIP followed by allele-specific qPCR in MCF7 cells and found that GATA3 was preferentially recruited to the *a*-allele of rs1391721 (**Figures 5G-H**). As further support, we investigated the correlation between *GATA3* and *TBX3* expression in the TCGA cohort. A stronger correlation was observed between *GATA3* and *TBX3* expression in normal breast as compared with the breast tumor samples (**Figure S6D**).

TBX3 is a T-Box TF that has been linked to tumorigenesis by impacting senescence and apoptosis as well as promoting proliferation and tumor formation^42^. To determine whether TBX3 can promote a tumorigenic phenotype in breast cells, we stably overexpressed or repressed TBX3 in the human mammary epithelial (HMLE) cell line and the MCF7 breast cancer cell line. HMLEs have been engineered to express h*TERT* and the SV40 large-T antigen and can grow in soft agar and form tumors in immune-deficient mice only upon introduction of an additional oncogenic insult^43^. Overexpression of TBX3 in HMLE cells resulted in a significant increase in cell colony growth in soft agar, suggesting that overexpression promotes anchorage-independent growth (**Figures 6A** and **S6E**), while CRISPR/Cas9-mediated TBX3 silencing showed a reciprocal effect (**Figures 6B**). These results are consistent with our *in vitro* data which indicated breast cancer risk was likely associated with increased TBX3 expression. The HMLE-TBX3 overexpressing cells were also injected into the mammary fat pads of nude mice, but no tumors were observed, suggesting elevated levels of TBX3 alone is not enough to promote tumor development from these cells. In contrast, overexpression of TBX3 in MCF7 cells decreased cell colony growth in soft agar (**Figures 6C** and **S6F**), while depletion of TBX3 by targeting dCas9-KRAB to the *TBX3* promoter resulted in a significant increase in growth (**Figures 6D** and **S6G**). To further investigate TBX3 in tumor growth, TBX3-depleted MCF7 cells were injected into the mammary fat pads of nude mice. Compared to control cells, reduced TBX3 levels resulted in a marked increase in tumor growth *in vivo* (**Figures 6E** and **6F**), which was reflected in increased tumor weights (**Figure 6G**). As reported previously ^44^, these data suggest that TBX3 can be oncogenic or tumor suppressive depending on cellular context.

**Figure 6.**
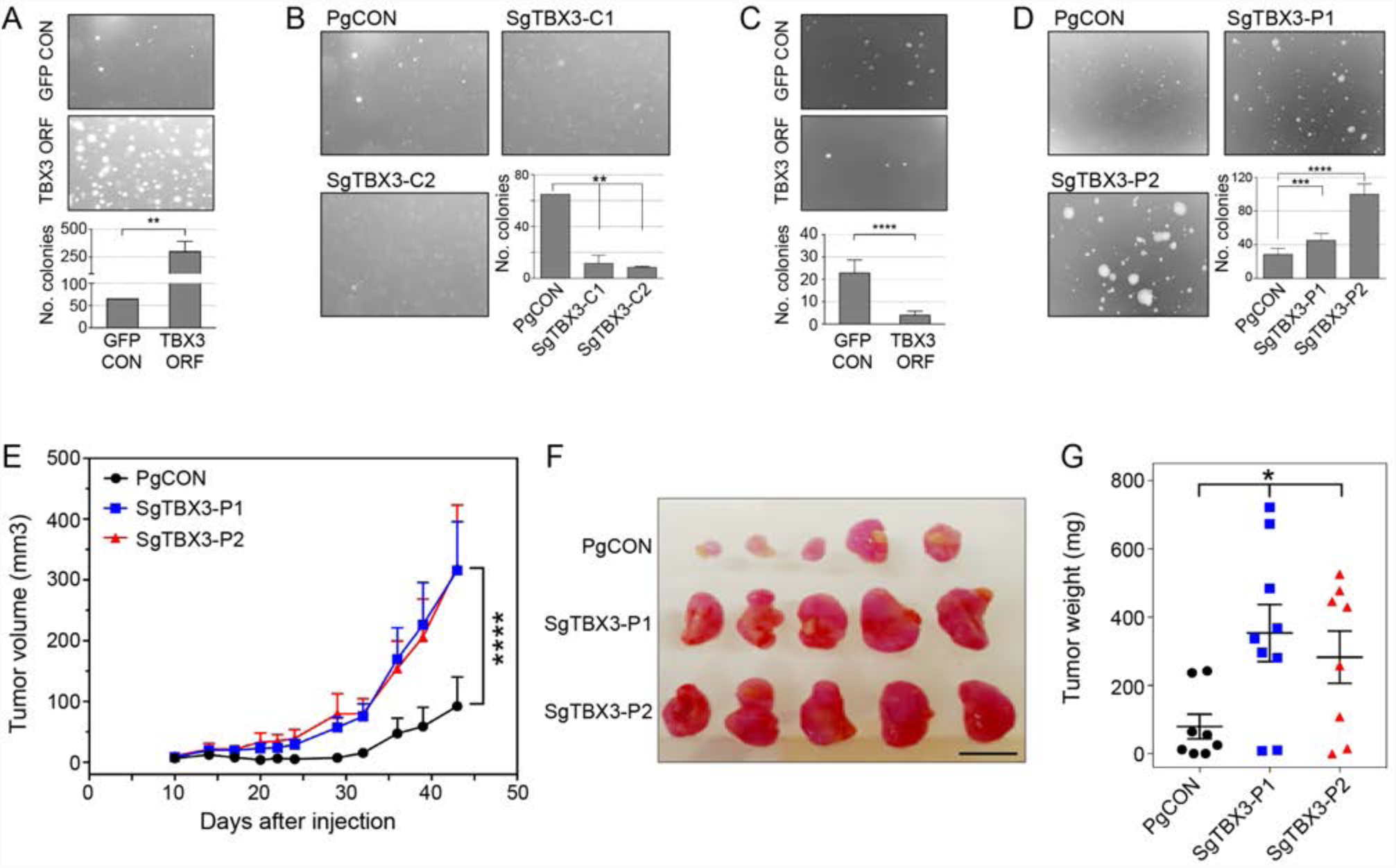
Altered TBX3 levels affect breast cell growth and tumor formation. **(a)** Representative images of colonies grown in soft agar for HMLE-control (GFP CON) and HMLE-TBX3 overexpressing cells (TBX3 ORF). The graph depicts the total number of HMLE colonies formed. Error bars represent the SEM (n = 2). **(b)** Representative images of colonies grown in soft agar for HMLE-control (PgCON) and HMLE-CRISPR/Cas9 TBX3 edited cells (SgTBX3-C1/C2). The graph depicts the total number of HMLE colonies formed. Error bars represent the SEM (n = 2). **(c)** Representative images of colonies grown in soft agar for MCF7-control (GFP CON) and MCF7-TBX3 overexpressing cells (TBX3 ORF). The graph depicts the total number of MCF7 colonies formed. Error bars represent the SEM (n = 4). **(d)** Representative images of colonies grown in soft agar for MCF7-control (PgCON) and MCF7-dCas9-KRAB TBX3 repressed cells (SgTBX3-P1/P2). The graph depicts the total number of MCF7 colonies formed. Error bars represent the SEM (n = 4). (**a-d**) p values were determined by two-way ANOVA followed by Dunnett’s multiple-comparison test (**p<0.01, ***p<0.001, ****p<0.0001). **(e)** MCF7-control (PgCON) or MCF7-dCas9-KRAB TBX3 repressed cells (SgTBX3-P1/P2) were injected into the mammary fat pads of nude mice. Tumor growth curves for each group are shown. Values are shown as average tumor volumes at each time point. Error bars represent the SEM (n = 8-9 mice per group). **(f)** Tumors of individual mice were dissected at day 44 post injection. The five largest tumors of each group are shown. The scale bar represents 1 cm. **(g)** Plot of the individual weights of tumors with mean and SEM shown by cross-bar and errors. **(e, g)** Mann–Whitney *U* test was used to compare differences between groups (*p<0.05, ****p<0.0001).

## DISCUSSION

The field of 3D chromatin interaction mapping is rapidly changing how we view the genome and is revealing important insights into disease biology. Interpretation of findings from GWAS has particularly benefited from the influx of chromatin data, allowing more accurate mapping and redefining of candidate causal genes. In this study, we generated high-resolution chromatin maps in human breast cells to delineate gene-regulatory interactions between breast cancer CCVs and target genes. We used two independent algorithms to score chromatin interactions. Peaky assisted identification of the probable direct contacts from long stretches of CHiCAGO-identified interactions.

This proved useful when examining PIRs as we were able to further prioritize the list of CCVs, which will be valuable in future in-depth functional studies. The de-prioritized variants may simply represent those in linkage disequilibrium with the true causal variant(s). Similarly, we observed an overlap between CHiCAGO-and Peaky-detected target genes, but noted that a proportion was detected by only one method. This was not unexpected given the different statistical models, and further studies will be required to establish parameters for improved resolution of direct interactions. Collectively, we could identify 651 candidate target genes at 139 independent breast cancer risk signals. Of particular interest for post-GWAS functional studies, 65 signals could be prioritized to one or two candidate target genes (**Table 1**). Some of the listed genes have functional data linking breast cancer CCVs to altered target gene expression, including *ESR1* ^45^, *FGFR2* ^46^ and *IGFBP5* ^47^, but most are still uncharacterized.

**Table 1.**
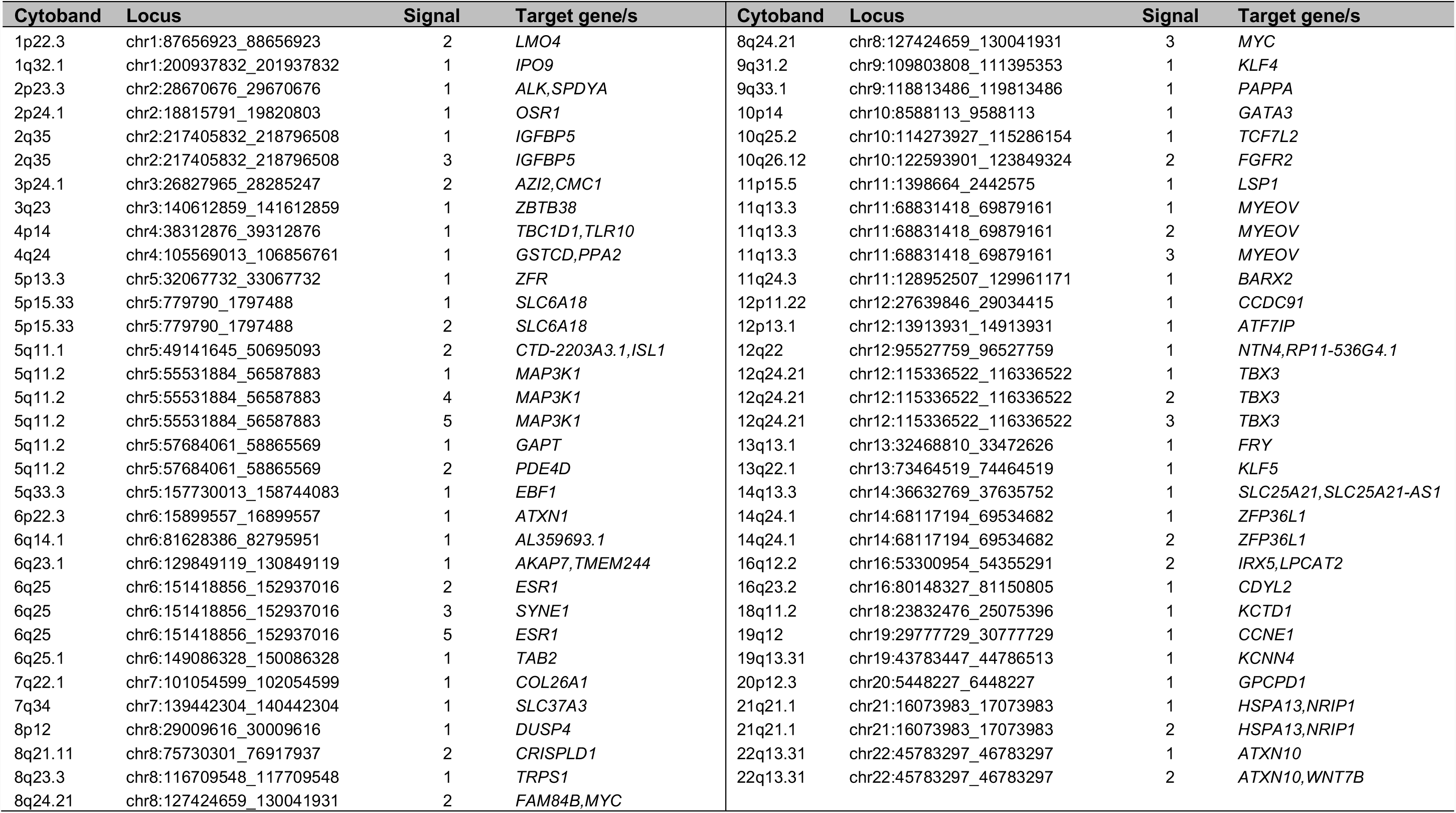
Independent breast cancer risk signals with ≤ two candidate protein-coding genes.

| Cytoband | Locus | Signal | Target gene/s | Cytoband | Locus | Signal | Target gene/s |
| --- | --- | --- | --- | --- | --- | --- | --- |
| 1p22.3 | chr1:87656923_88656923 | 2 | *LMO4* | 8q24.21 | chr8:127424659_130041931 | 3 | *MYC* |
| 1q32.1 | chr1:200937832_201937832 | 1 | *IPO9* | 9q31.2 | chr9:109803808_111395353 | 1 | *KLF4* |
| 2p23.3 | chr2:28670676_29670676 | 1 | *ALK,SPDYA* | 9q33.1 | chr9:118813486_119813486 | 1 | *PAPPA* |
| 2p24.1 | chr2:18815791_19820803 | 1 | *OSR1* | 10p14 | chr10:8588113_9588113 | 1 | *GATA3* |
| 2q35 | chr2:217405832_218796508 | 1 | *IGFBP5* | 10q25.2 | chr10:114273927_115286154 | 1 | *TCF7L2* |
| 2q35 | chr2:217405832_218796508 | 3 | *IGFBP5* | 10q26.12 | chr10:122593901_123849324 | 2 | *FGFR2* |
| 3p24.1 | chr3:26827965_28285247 | 2 | *AZI2,CMC1* | 11p15.5 | chr11:1398664_2442575 | 1 | *LSP1* |
| 3q23 | chr3:140612859_141612859 | 1 | *ZBTB38* | 11q13.3 | chr11:68831418_69879161 | 1 | *MYEOV* |
| 4p14 | chr4:38312876_39312876 | 1 | *TBC1D1,TLR10* | 11q13.3 | chr11:68831418_69879161 | 2 | *MYEOV* |
| 4q24 | chr4:105569013_106856761 | 1 | *GSTCD,PPA2* | 11q13.3 | chr11:68831418_69879161 | 3 | *MYEOV* |
| 5p13.3 | chr5:32067732_33067732 | 1 | *ZFR* | 11q24.3 | chr11:128952507_129961171 | 1 | *BARX2* |
| 5p15.33 | chr5:779790_1797488 | 1 | *SLC6A18* | 12p11.22 | chr12:27639846_29034415 | 1 | *CCDC91* |
| 5p15.33 | chr5:779790_1797488 | 2 | *SLC6A18* | 12p13.1 | chr12:13913931_14913931 | 1 | *ATF7IP* |
| 5q11.1 | chr5:49141645_50695093 | 2 | *CTD-2203A3.1,ISL1* | 12q22 | chr12:95527759_96527759 | 1 | *NTN4,RP11-536G4.1* |
| 5q11.2 | chr5:55531884_56587883 | 1 | *MAP3K1* | 12q24.21 | chr12:115336522_116336522 | 1 | *TBX3* |
| 5q11.2 | chr5:55531884_56587883 | 4 | *MAP3K1* | 12q24.21 | chr12:115336522_116336522 | 2 | *TBX3* |
| 5q11.2 | chr5:55531884_56587883 | 5 | *MAP3K1* | 12q24.21 | chr12:115336522_116336522 | 3 | *TBX3* |
| 5q11.2 | chr5:57684061_58865569 | 1 | *GAPT* | 13q13.1 | chr13:32468810_33472626 | 1 | *FRY* |
| 5q11.2 | chr5:57684061_58865569 | 2 | *PDE4D* | 13q22.1 | chr13:73464519_74464519 | 1 | *KLF5* |
| 5q33.3 | chr5:157730013_158744083 | 1 | *EBF1* | 14q13.3 | chr14:36632769_37635752 | 1 | *SLC25A21,SLC25A21-AS1* |
| 6p22.3 | chr6:15899557_16899557 | 1 | *ATXN1* | 14q24.1 | chr14:68117194_69534682 | 1 | *ZFP36L1* |
| 6q14.1 | chr6:81628386_82795951 | 1 | *AL359693.1* | 14q24.1 | chr14:68117194_69534682 | 2 | *ZFP36L1* |
| 6q23.1 | chr6:129849119_130849119 | 1 | *AKAP7,TMEM244* | 16q12.2 | chr16:53300954_54355291 | 2 | *IRX5,LPCAT2* |
| 6q25 | chr6:151418856_152937016 | 2 | *ESR1* | 16q23.2 | chr16:80148327_81150805 | 1 | *CDYL2* |
| 6q25 | chr6:151418856_152937016 | 3 | *SYNE1* | 18q11.2 | chr18:23832476_25075396 | 1 | *KCTD1* |
| 6q25 | chr6:151418856_152937016 | 5 | *ESR1* | 19q12 | chr19:29777729_30777729 | 1 | *CCNE1* |
| 6q25.1 | chr6:149086328_150086328 | 1 | *TAB2* | 19q13.31 | chr19:43783447_44786513 | 1 | *KCNN4* |
| 7q22.1 | chr7:101054599_102054599 | 1 | *COL26A1* | 20p12.3 | chr20:5448227_6448227 | 1 | *GPCPD1* |
| 7q34 | chr7:139442304_140442304 | 1 | *SLC37A3* | 21q21.1 | chr21:16073983_17073983 | 1 | *HSPA13,NRIP1* |
| 8p12 | chr8:29009616_30009616 | 1 | *DUSP4* | 21q21.1 | chr21:16073983_17073983 | 2 | *HSPA13,NRIP1* |
| 8q21.11 | chr8:75730301_76917937 | 2 | *CRISPLD1* | 22q13.31 | chr22:45783297_46783297 | 1 | *ATXN10* |
| 8q23.3 | chr8:116709548_117709548 | 1 | *TRPS1* | 22q13.31 | chr22:45783297_46783297 | 2 | *ATXN10,WNT7B* |
| 8q24.21 | chr8:127424659_130041931 | 2 | *FAM84B,MYC* |  |  |  |  |

A recent study used CHi-C to identify 110 putative target genes at 33 breast cancer risk loci ^48^. Surprisingly, only 30 of the 110 genes were also identified in our study. The lack of concordance may firstly result from a fundamental difference in capture design; Baxter et al. was based on SNPs correlated with the published SNP (r^2^?0.2); whereas the present study captures only those fragments containing CCVs based on fine-mapping analysis of a very large association dataset. In addition, the design used by Baxter and colleagues included many examples where oligonucleotide probes were tiled across large genomic regions rather than restricted to individual *HindIII* fragments. Baxter et al. also reported multiple genes as putative targets at some risk signals, while our analysis of the same signals prioritized only one or two genes. For example, at 11p15.5 Baxter *et al.* identified nine target genes, whereas our combined statistical analyses reduced this number to just two candidates, *LSP1* and *MIR4298*.

We acknowledge that some CCV-target gene interactions may have been missed due to intrinsic biases in the capture. False negatives may result from lack of suitable baits for some CCV-and promoter-containing fragments, short range contact constraints or due to the transient and cell type-specific nature of regulatory chromatin interactions. It is also important to keep in mind that interactions between a CCV and gene promoter do not infer causality. It is likely that correlated CCVs within some signals have no effect on TF binding or enhancer activity, or they may act via alternate mechanisms. Consistent with other GWAS follow-up studies ^49^, our results support the hypothesis that *cis*-acting regulatory variation is a predominant molecular mechanism at breast cancer risk signals. However, we saw no CCV-target gene looping interactions at 57 (out of 196) risk signals.

Twelve signals contained promoter or coding CCVs, suggesting that direct gene alteration is a probable mechanism underlying these risk associations. The remaining signals (n=45) contained baited variant or promoter fragments, but the lack of detected CCV-gene interactions suggests mechanisms other than distal regulation. A recent study has incorporated some of the proposed alternate CCV mechanisms together with the distally-regulated genes from this study to generate a complete catalog of candidate target genes and biological pathways ^5^.

We provided functional evidence that breast cancer risk at 12q24 is driven by the TF, TBX3. TBX3 is overexpressed in many cancers including breast cancer, and contributes to oncogenesis at multiple levels including promotion of proliferation, tumor formation and metastasis ^42^. Consistent with previous findings, our *in vitro* data indicate that the signal 1 CCVs likely act to increase TBX3 expression through recruitment of GATA3 to the CCV site, resulting in increased looping of the risk CCV-containing enhancer to the *TBX3* promoter. Several studies have suggested that TBX3 may also function as a tumor suppressor depending on the cellular context ^44^. Indeed, in MCF7 breast cancer cells, we showed that TBX3 repression promoted colony formation and *in vivo* tumor formation. Furthermore, somatic *TBX3* mutations in primary breast tumors are predominantly loss-of-function through impaired transcriptional repression ^50^. Interestingly, a recent report showed that many of these “double agent” genes are TFs and that breast cancer is the second most common cancer type associated with dual-function genes ^51^. The molecular mechanisms underlying this duality are largely unknown, but differing mutation spectrums, interaction partners and cellular contexts have been implicated. Dual-function genes likely contribute to the heterogeneity of cancer cells and some are already considered promising targets for breast cancer therapy. It will therefore be important to refine therapeutic strategies to selectively block one function without compromising the other.

In summary, we report the most comprehensive study linking regulatory CCVs to candidate breast cancer genes. This forms an important resource for the breast cancer research community that will facilitate generation of hypotheses, functional experimentation as well as insights into breast cancer biology. We anticipate that many of the candidate target genes may represent drug repositioning opportunities or be suitable for future drug targeting.

## METHODS

### Data availability

Raw sequencing data has been deposited at EBI: PRJEB29716. Processed Capture Hi-C data is available from https://osf.io/2cnw7/. Processed chromatin interaction data can be visualized at the Washington Epigenome Browser via https://bit.ly/2rnCqS8.

### Code availability

The custom scripts used during the study are available from the corresponding author on reasonable request.

### URLs

HiCUP, http://bioinformatics.babraham.ac.uk/projects/hicup/overview; CHiCAGO, http://regulatorygenomicsgroup.org/chicago; Peaky, http://github.com/cqgd/pky; Integrative Genomics viewer, http://software.broadinstitute.org/software/igv; Graphpad Prism, http://graphpad.com/scientific-software/prism/; Cutadapt (version 1.9), http://pypi.python.org/pypi/cutadapy/19.1; BWA-MEM (version 0.7.12), http://bio-bwa.sourceforge.net; Samtools (version 1.1), http://htslib.org; Picard (version 1.129), http://github.com/broadinstitute/picard; qProfiler, http://sourceforge.net/p/adamajava/wiki/qProfiler/; MACS2, http://github.com/taoliu/MACS; HOMER, http://homer.ucsd.edu/homer/; R, https://www.r-project.org; JASPAR, http://jaspar.genereg.net/.

### Cell lines

Estrogen receptor positive (ER+) breast cancer cell lines MCF7 and T47D were grown in RPMI medium with 10% (vol/vol) fetal bovine serum (FBS), 1 mM sodium pyruvate, 10 ?g/ml insulin, and 1% (vol/vol) antibiotics. ER-breast cancer cell lines MDAMB231 and Hs578T were grown in DMEM medium with 10% (vol/vol) FBS and 1% (vol/vol) antibiotics. The B80T5 mammary epithelial cell line (provided by Roger Reddel, CMRI, Australia) was grown in RPMI medium with 10% (vol/vol) FBS and 1% (vol/vol) antibiotics. The MCF10A mammary epithelial cell line was grown in DMEM/F12 medium with 5% (vol/vol) horse serum, 10 μ?g/ml insulin, 0.5 μ?g/ml hydrocortisone, 20 ng/ml epidermal growth factor, 100 ng/ml cholera toxin, and 1% (vol/vol) antibiotics. Cell lines were maintained under standard conditions (37 °C, 5% CO_2_), tested for *Mycoplasma* and profiled for short tandem repeats.

### Hi-C library preparation

Hi-C libraries were prepared from 4-8x10^7^ cells per library (two biological replicates per cell line; three replicates for the T47D VCHi-C) as described previously^11^, but using in-nucleus ligation as described in ^52^. The immobilized Hi-C libraries were amplified using the SureSelect^XT^ ILM Indexing pre-capture primers (Agilent Technologies) with eight PCR amplification cycles. Each Hi-C library (750 ng) was hybridized and captured individually using the SureSelect^XT^ Target Enrichment System reagents and protocol (Agilent Technologies). After library enrichment, a post-capture PCR amplification step was carried out using SureSelect^XT^ ILM Indexing post-capture primers (Agilent Technologies) with 14-16 PCR amplification cycles.

### Biotinylated RNA bait library design

The SureSelect^XT^ Custom Target Enrichment Arrays were designed using the eARRAY software (Agilent Technologies). For the VCHi-C, biotinylated 120-mer RNA baits were designed to both ends of *HindIII* restriction fragments that contained at least one CCV^5^. A total of 1448 *HindIII* fragments were captured, covering 6044/7394 CCVs. For the PCHi-C, biotinylated 120-mer RNA baits were designed to both ends of *HindIII* restriction fragments that overlapped annotated promoters within 1 Mb of CCVs ^5^. A total of 4049 *HindIII* fragments were captured, overlapping 2298 Ensembl-annotated promoters (GRCh38) ^16^. A bait sequence was accepted if its GC content was between 25-65%, the sequence contained no more than two consecutive nucleotides of the same identity, and was within 330 bp of the *HindIII* restriction fragment end. Repetitive elements were masked using SureDesign masking tools with the highest level of stringency.

### Sequencing of CHi-C libraries

PCHi-C and VCHi-C libraries were sequenced on the Illumina HiSeq 2500 platform (Kinghorn Centre for Clinical Genomics, Australia). Two PCHi-C or three VCHi-C libraries were multiplexed per sequencing lane.

### PCHi-C and VCHi-C sequence alignment and data processing

Raw sequencing reads were truncated, mapped to the hg19 reference genome, and filtered using the HiCUP pipeline ^53^. Individual library statistics are presented in **Table S1**. Significant interactions were identified using the CHiCAGO pipeline^23^. For both captures, replicate libraries for each cell line were analyzed separately to learn weights which were then used to merge replicates into a single dataset per cell type. Interactions with CHICAGO scores ≥ 5 in at least one cell type were considered high-confidence interactions.

### Principal component and cluster analyses

Principal Component Analysis (PCA) of the CHiCAGO interaction scores was performed for both variant and promoter capture arrays for each individual biological replicate. Interaction length <2 Mb and CHiCAGO score >0 were included. PCA was performed using the R utility *prcomp* with unit variance scaling. Hierarchical clustering with average linkage based on Euclidian distances was performed on the 1000 interactions with most variance using R’s *heatmap.2* function. Cell types were clustered based on profiles including interactions with CHiCAGO score >=5 and length <2 Mb. Interactions with score >=5 in at least one cell line were considered.

### PCHi-C and VCHi-C concordance

To examine the overall concordance between promoter and variant captures, we identified interactions common to both experiments from the full range of CHiCAGO scores (>0) for each cell type. The Pearson correlation between CHiCAGO scores for interactions from each of the captures was computed. Interactions scores for each capture were plotted after inverse hyperbolic sine (asinh) transformation with loess smoothed regression lines.

### Enrichment of genomic features within interacting regions

Positions of genomic features including DNase-seq peak, histone modification ChIP-seq peaks, transcription factor ChIP-seq peaks (web links provided in **Table S7**) and ATAC-seq peaks were intersected with PIRs from each cell line. Enrichment was estimated by comparing to a set of background PIRs generated by maintaining the distribution of interaction distances and interaction counts relative to promoter baits for each cell type. Interactions were grouped in 50 kb distance bins, and 100 sets of random PIR sets were built for each cell line. We removed baited fragments from the pool of possible PIRs. Z scores were calculated for each genomic annotation.

### Fine-mapping of chromatin contacts

PCHi-C and VCHi-C contact mapping was performed using the *Peaky* Bioconductor package ^30^. We first pooled aligned reads from replicate CHi-C libraries. Probable interaction-driving contacts were then modelled for each bait from each cell line independently. We maintained the default ? value (5) for each bait. Two parallel chains were run and correlation between MPPC values for interacting prey fragments were tested until r > 0.75 (typically after 20^6^ iterations). We achieved successful convergence for >93% tested baits. Distributions derived from parallel chains were then merged to generate cell-type and bait-specific contact maps. An arbitrary MPPC threshold of 0.1 was used for downstream analysis.

### Expression quantitative trait loci analysis

To determine whether eSNP-target gene pairs were over-represented within captured interactions, we assigned interactions to random promoters within the same chromosome. This randomization procedure was repeated 10,000 times. The frequency of eSNP-gene occurrences within interactions was then tallied in the observed interaction set and compared to random expectation.

### ATAC-seq library preparation and data analysis

ATAC-seq was performed as previously described^54^. Briefly, 5x10^4^ cells were resuspended in lysis buffer (10 mM Tris-HCl (pH 7.4), 10 mM NaCl, 3 mM MgCl_2_, 0.1% (vol/vol) IGEPAL CA-630), then centrifuged at 5000xg for 10 min at 4°C. Pellets were resuspended in TD buffer (10 mM Tris (pH 7.6), 5 mM MgCl_2_, 10% (vol/vol) dimethylformamide) and 2.5 ?l of TDE1 enzyme (Illumina). Transposed fragments were purified using a MinElute PCR purification kit (QIAGEN), then amplified and indexed with unique library indices using NEBNext High-Fidelity 2x PCR Master Mix (New England BioLabs). PCR products were purified with AMPure XP beads (Beckman-Coulter) and quantified with a Qubit dsDNA High-Sensitivity Assay kit (Thermo Fisher Scientific) and BioAnalyzer High Sensitivity DNA Kit (Agilent Technologies). Pools of six libraries were sequenced per lane on an Illumina HiSeq 2500 (Kinghorn Centre for Clinical Genomics). Raw sequencing reads were trimmed for adapter sequences using Cutadapt (version 1.9;^55^) and aligned using BWA-MEM (version 0.7.12;^56^) to the GRCh37 assembly. The aligned reads were coordinate sorted using Samtools (version 1.1;^57^) and duplicate alignments were marked with Picard (version 1.129). Qprofiler assessed the sequence quality and provide fragment length distribution. Peaks were called for each sample using MACS2^58^. Peak annotation was performed using HOMER^59^.

### 3C validation

3C libraries were generated using *HindIII* as described previously^47^. 3C interactions were quantified by real-time PCR (qPCR) using primers designed within restriction fragments (**Table S6**). qPCR was performed on a RotorGene 6000 using MyTaq HS DNA polymerase (Bioline) with the addition of 25?M Syto9, annealing temperature of 66°C and extension time of 30s. 3C analyses were performed in two independent 3C libraries from each cell line quantified in duplicate. BAC clones covering each region were used to create artificial libraries of ligation products to normalize for PCR efficiency. Data were normalized to the signal from the BAC clone library and, between cell lines, by reference to a region within *GAPDH*.

### CRISPR/Cas9 interference and cutting

For CRISPR interference (CRISPRi), the sgRNA targets (listed in **Table S6**), Cas9 binding handle and terminator sequences were synthesized (Integrated DNA Technologies, IDT) and cloned into the lentiviral vector pgRNA-humanized. Virus-like particles (VLPs) containing either dCas9-KRAB or a targeting sgRNA were generated by transfection of HEK293 cells with Lipofectamine 2000 (Thermo Fisher Scientific). Cells were cotransfected with the packaging plasmid pCMV-dR8.91, the VSV-G envelope expression plasmid pCMV-VSV-G, and with either pHR-SFFV-dCas9-BFP-KRAB or pgRNA-humanized. VLPs were collected from culture supernatants, mixed in equal volume, and transduced into MCF7 cells. Cells expressing both mCherry (via pgRNA-humanized) and blue fluorescent protein (via dCas9-KRAB) were isolated by FACS on an ARIA IIIu (Becton-Dickinson). For CRISPR cutting (CRISPRc), the GFP control and sgRNA targets (listed in **Table S6**) were synthesized (IDT) and cloned into the pXPR_011 lentiviral vector. Virus-like particles (VLPs) containing the GFP control or targeting sgRNAs were generated by transfection of HEK293 cells with FuGene (Promega). VLPs were collected from culture supernatant, transduced into HMLE-Cas9 cells, and selected using puromycin for at least 48 h.

### Quantitative real-time PCR (qPCR)

Complementary DNA (cDNA) was synthesized from RNA samples using SuperScript III (Invitrogen). qPCR was performed using TaqMan assays (Thermo Fisher Scientific; listed in **Table S6**).

### Plasmid construction and reporter assays

The *TBX3* promoter-driven luciferase construct was generated by insertion of a PCR amplified promoter fragment into the NheI and *HindIII* sites of the pGL3-basic vector (primers are listed in **Table S6**). The 12q24 signal 1 enhancer, containing either the risk or protective CCV alleles, was synthesized as gBlocks (IDT) and cloned into the BamHI and SalI sites of the *TBX3*-promoter construct (coordinates are listed in **Table S6**). Sanger sequencing of all constructs confirmed variant incorporation. MCF7 cells were transfected with equimolar amounts of luciferase reporter plasmids and pRL-TK transfection control plasmid with Lipofectamine 3000 (Thermo Fisher Scientific).Luciferase activity was measured 24 h post-transfection by the Dual-Glo Luciferase Assay System (Promega). To correct for any differences in transfection efficiency or cell lysate preparation, *Firefly* luciferase activity was normalized to *Renilla* luciferase activity, and the activity of each construct was expressed relative to the reference promoter constructs, which were defined to have an activity of 1.

### Electromobility shift assays (EMSAs)

Gel shift assays were performed with MCF7 or BT474 nuclear lysates and biotinylated oligonucleotide duplexes (listed in **Table S6**). Nuclear lysates were prepared using the NE-PER nuclear and cytoplasmic extraction reagents (Thermo Fisher Scientific) as per the manufacturer’s instructions. Total protein concentrations in nuclear lysates were determined by Bradford’s method. Duplexes were prepared by combining sense and antisense oligonucleotides in NEBuffer2 (New England Biolabs) and heat annealing at 80°C for 10 min followed by slow cooling to 25°C for 1 h. Binding reactions were performed in binding buffer (10% (vol/vol) glycerol, 20 mM HEPES (pH 7.4), 1 mM DTT, protease inhibitor cocktail (Roche), 0.75 μg poly(dI:dC) (Sigma-Aldrich)) with 7.5 μg of nuclear lysate. For competition assays, binding reactions were pre-incubated with 1 pmol of competitor duplex (competitor sequences are listed in **Table S6**) at 25°C for 10 min before the addition of 10 fmol of biotinylated duplex and incubation at 25°C for 15 min. Reactions were separated on 10% (wt/vol) Tris-Borate-EDTA (TBE) polyacrylamide gels (Bio-Rad) in TBE buffer at 160 V for 40 min. Duplex-bound complexes were transferred onto Zeta-Probe positively-charged nylon membranes (Bio-Rad) by semi-dry transfer at 25 V for 20 min, then cross-linked onto the membranes under 254 nm ultra-violet light for 10 min. Membranes were processed with the LightShift Chemiluminescent EMSA kit (Thermo Fisher Scientific) as per the manufacturer’s instructions. Chemiluminescent signals were visualized with the C-DiGit blot scanner (LI-COR).

### Chromatin immunoprecipitation (ChIP)

MCF7 cells were cross-linked with 1% (wt/vol) formaldehyde at 37°C for 10 min, rinsed once with ice-cold PBS containing 5% (wt/vol) BSA and once with PBS, and harvested in PBS containing protease inhibitor cocktail (Roche). Harvested cells were centrifuged for 2 min at 3000 rpm. Cell pellets were resuspended in 0.35 ml of lysis buffer (1% (wt/vol) SDS, 10 mM EDTA, 50 mM Tris-HCl (pH 8.1)),protease inhibitor cocktail and sonicated three times for 15 s at 70% duty cycle (Branson SLPt) followed by centrifugation at 13000 rpm for 15 min. Supernatants were collected and diluted in dilution buffer (1% (wt/vol) Triton X-100, 2 mM EDTA, 150 mM NaCl, 20 mM Tris-HCl (pH 8.1)). Two micrograms of anti-GATA3 antibody (Santa Cruz) or control IgG (Santa Cruz) was prebound for 6 h to protein G Dynabeads (Thermo Fisher Scientific) and then added to the diluted chromatin for overnight immunoprecipitation. The magnetic bead-chromatin complexes were collected and washed six times in RIPA buffer (50 mM HEPES (pH 7.6), 1 mM EDTA, 0.7% (vol/vol) sodium deoxycholate, 1% (vol/vol) NP-40, 0.5 M LiCl), then twice with TE buffer. To reverse cross-linking, the magnetic bead complexes were incubated overnight at 65°C in elution buffer (1% (wt/vol) SDS, 0.1 M NaHCO_3_). DNA fragments were purified using the QIAquick Spin Kit (QIAGEN). For qPCR (primers are listed in **Table S6**), 2 ?l from a 100 ?l immunoprecipitated chromatin extraction were amplified for 40 cycles. All PCR products were sequenced by Sanger sequencing.

### TBX3 overexpression

The TBX3 overexpression construct (pLX307/TBX3) was generated by Gateway cloning from pDONR201 containing the full length TBX3 cDNA into the pLEX_307 lentiviral destination vector (Thermo Fisher Scientific). A negative control construct (pLX307/CON) was generated by excising TBX3 via *Nhe*I and *Spe*I restriction enzyme digestion and self-ligating the vector backbone. VLPs were generated from HEK293 cells transfected with pLX307/CON or pLX307/TBX3 as described above and transduced into HMLE or MCF7 cells. Transductants were selected with puromycin for at least 48 h.

### Western blotting

Cell pellets were lysed in RIPA buffer (50 mM Tris-HCl (pH 8.0), 150 mM NaCl; 1% (vol/vol) IGEPAL CA-630, 0.5% (vol/vol) sodium deoxycholate, 0.1% (wt/vol) SDS, 1 mM DTT, protease inhibitor cocktail) and clarified by centrifugation to remove cell debris. Forty micrograms of lysate supernatants were separated by SDS-polyacrylamide gel electrophoresis, electroblotted onto PVDF membranes by semi-dry transfer (Bio-Rad) and blocked in blocking buffer (1% (wt/vol) casein, 0.1% (vol/vol) Tween 20, PBS). TBX3 was detected with 1 μg/ml rabbit anti-TBX3 antibody (Thermo Fisher Scientific) and actin with 400 ng/ml of rabbit anti-actin antibody (Sigma-Aldrich). Primary antibodies were detected with horseradish peroxidase-conjugated goat anti-rabbit IgG (Cell Signaling). Detected proteins were visualized with enhanced chemiluminescence substrate (Bio-Rad) and the G:BOX Chemi XX6 gel documentation system (Syngene).

### Soft agar colony formation assay

Six-well plates were layered with 0.6% (wt/vol) noble agar (Becton-Dickinson) in RPMI or DMEM medium supplemented with 10% (vol/vol) FBS and antibiotics and allowed to set at 4°C. Twenty-four hours later, cells were trypsinized and 8x10^3^ MCF7 or 5x10^4^ HMLE cells were resuspended in 0.3% (wt/vol) noble agar and plated on top of bottom agar layers (3 wells/cell line). Colonies were imaged after 3-4 weeks using a Leica MZ FLIII stereo microscope.

### Cell proliferation assay

Cell proliferation was measured using a label-free, non-invasive cellular confluence assay on the IncuCyte Live-Cell Imaging System (Essen Bioscience). MCF7 cells were seeded at 20,000 cells/well into 24-well plates and imaged on the IncuCyte using a 10x objective lens every 3 h over 7 days. Imaging was performed in an incubator maintained at 37°C under a 5% CO_2_ atmosphere. Cell confluence in each well was measured using IncuCyte ZOOM 2016A software, and the data analyzed using GraphPad Prism.

### Mouse tumor xenograft model

A cholesterol-based pellet containing 17β-estradiol (0.72 mg, 90-day slow release, Innovative Research of America) was implanted subcutaneously in the interscapular region of 8-week old female BALB/c-Foxn1^nu^/Arc mice. Three days later, MCF7 CRISPRi-suppressed cells (1x10^7^ cells/mouse) were injected into the 4^th^ right mammary fatpad (8-9 mice per cell line). Tumor volumes were measured with a digital caliper every second day until the experimental end stage approved by the QIMR Berghofer animal ethics committee; 525 mm^3^ according to the formula (π×length×width^2^/6).

## ACKNOWLEDGEMENTS

This work was supported by grants from the National Health and Medical Research Council of Australia (NHMRC; 1058415 and 1120563), Cancer Council Queensland (1099810) and Perpetual IMPACT Program (IPAP2017/1497). S.L.E. and N.W. are NHMRC Senior Research Fellows (1135932 and 1139071). G.C.T is an NHMRC Senior Principle Research Fellow (1117073). J.D.F was supported by a Fellowship from the National Breast Cancer Foundation of Australia. N.A. was co-funded by a QIMR Berghofer International PhD Scholarship and a University of Queensland Research Training Scholarship. This project has received funding from the European Union’s Horizon 2020 Marie Skłodowska-Curie Individual Fellowships programme under grant agreement No [MSCA-IF-2014-EF-656144]. The results published here are in part based upon data generated by the TCGA Research Network. The authors declare no competing financial interests.

## AUTHOR CONTRIBUTIONS

Conceptualization, J.D.F and S.L.E. Bioinformatic and statistical analyses: J.B., M.M.M., P.M., S.K. K.M., D.R.B., L.F. Functional analyses: H.S., L.G.L, N.T., K.M.H., S.Kaufmann, N.H., S.H., J.S.L.,

K.N. Supervision, A.C.A., A.M.D., D.F.E., N.W., J.R., A.M., G.C.T. Writing, J.B., J.D.F. and S.L.E., with contributions from all authors.

---

**Figure S1.**
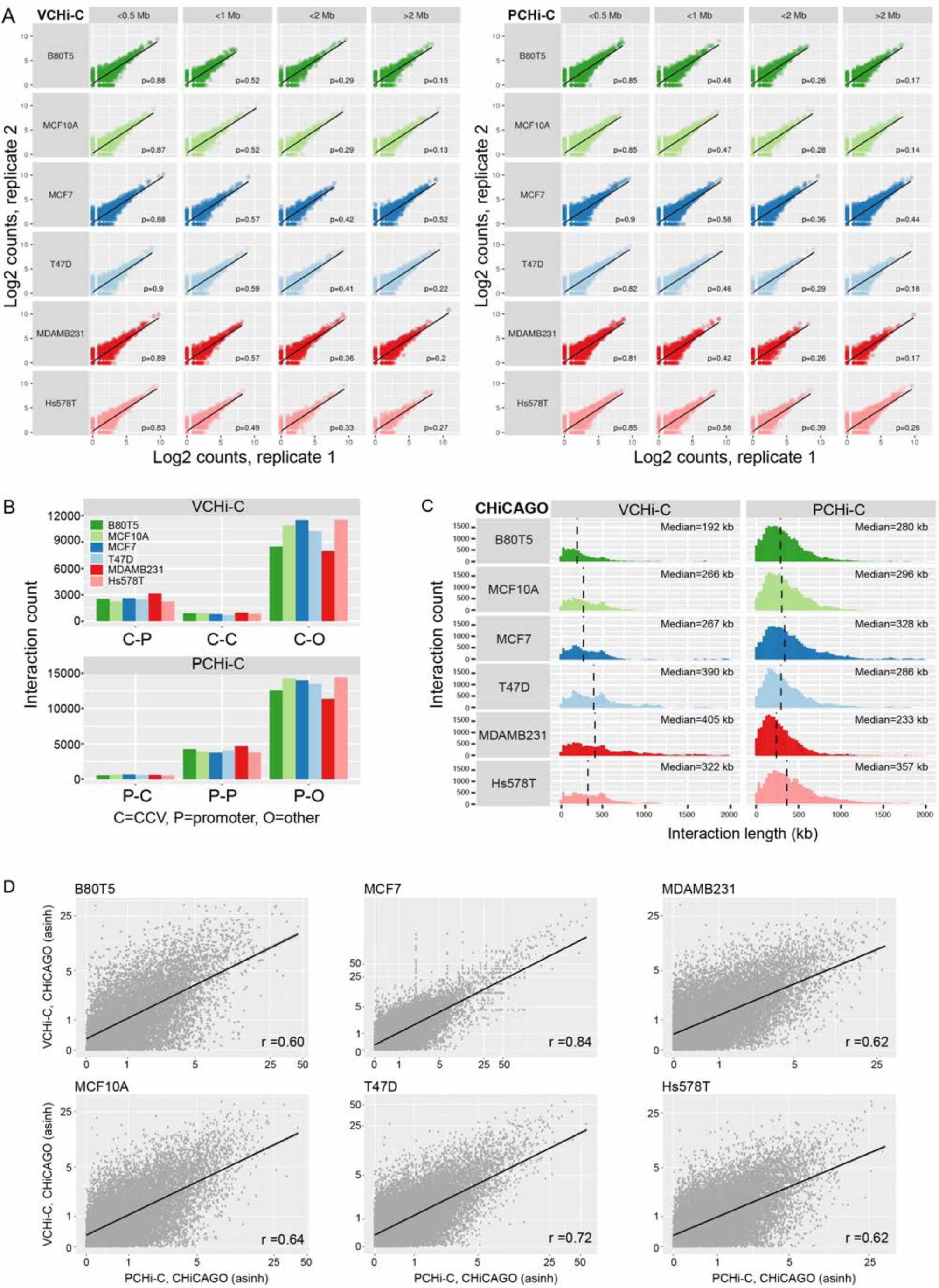
VCHi-C and PCHi-C CHiCAGO-identified interaction characteristics, related to Figure 1. **(A)** Scatter plots showing the correlation between duplicate VCHi-C or PCHi-C libraries based on the number of raw di-tags mapping to interaction fragment pairs. The analysis was stratified by cell line (rows) and distance between interacting fragments (columns). ρ is Spearman’s correlation; the black lines represent the linear regression fit. **(B)** The abundance of different classes of CHiCAGO-scored VCHi-C (upper panel) and PCHi-C (lower panel) interactions. **(C)** Distribution of CHiCAGO-scored interaction lengths in each breast cell line. Dashed black vertical lines denote the median interaction length. **(D)** Scatter plots showing the concordance of inverse hyperbolic sine (asinh)-transformed CHiCAGO-scored VCHi-C versus PCHi-C interactions in the respective breast cell lines. r is Pearson’s correlation; the black lines represent the linear regression fit.

**Figure S2.**
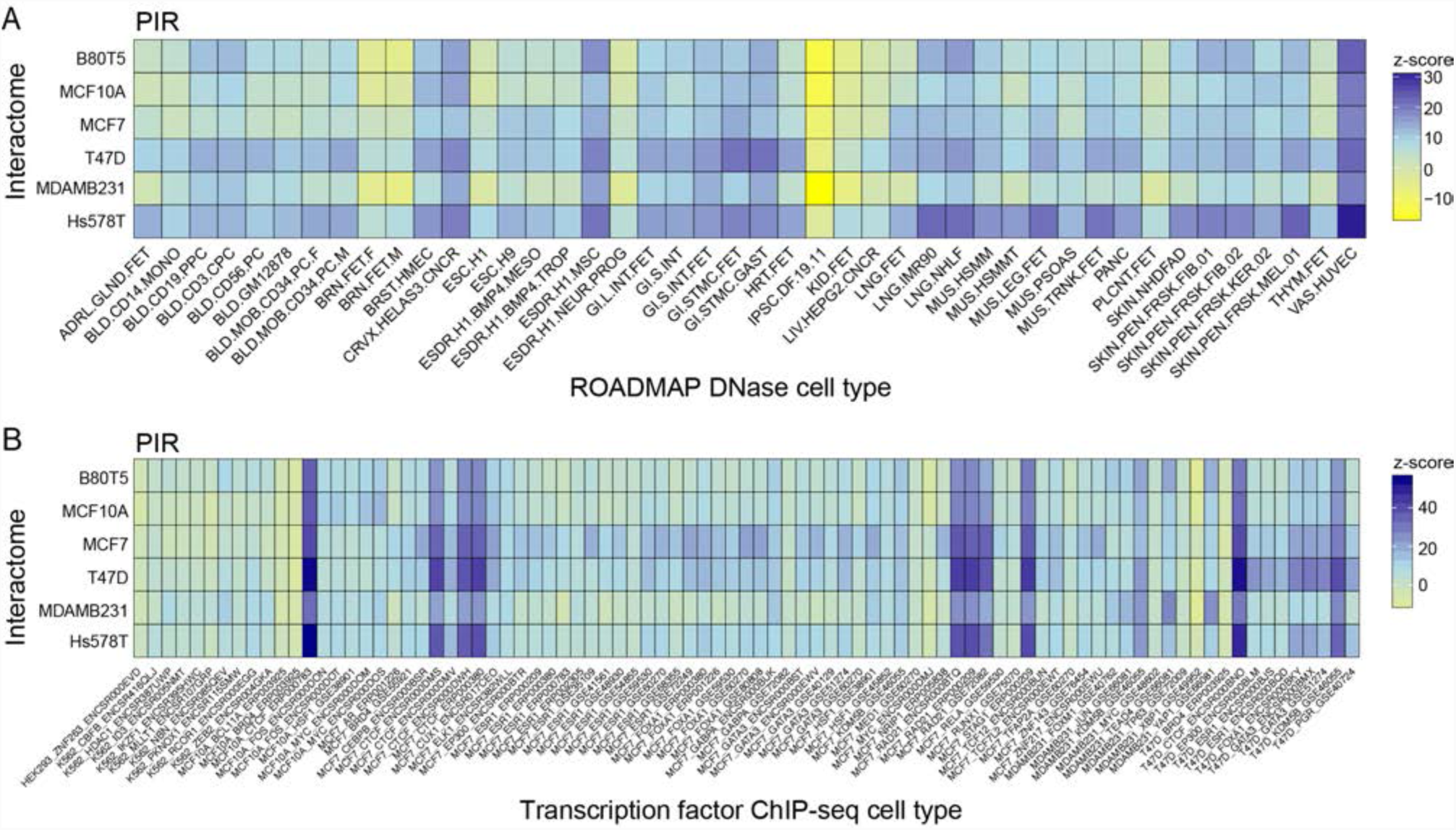
PIRs are enriched for breast-specific regulatory features, related to Figure 2. **(A)** Heatmap showing promoter-interacting region (PIR) enrichment for DNase I hypersensitivity sites in additional ROADMAP cell types, expressed as z-scores. **(B)** Heatmap showing PIR enrichment for transcription factor binding in breast (additional datasets) and other cell types, expressed as z-scores.

**Figure S3.**
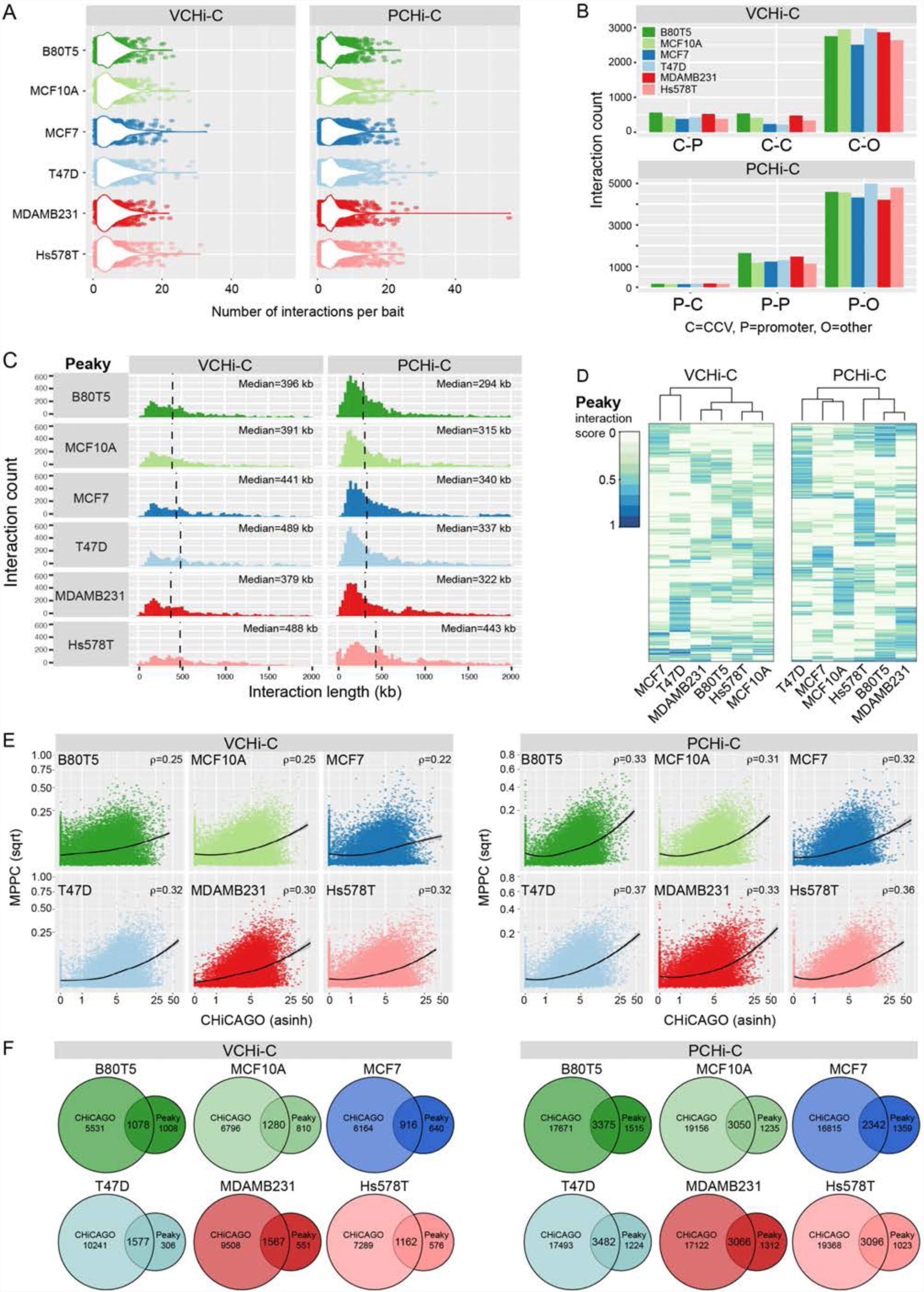
VCHi-C and PCHi-C Peaky-identified interaction characteristics, related to Figures 3 and 4. **(A)** Distribution of Peaky-scored interaction number per bait per cell line (combined biological replicates). **(B)** The abundance of different classes of Peaky-scored VCHi-C (upper panel) and PCHi-C (lower panel) interactions. **(C)** Distribution of Peaky-scored interaction lengths in each breast cell line. Dashed black vertical lines denote the median interaction length. **(D)** Agglomerative hierarchical clustering for the VCHi-C and PCHi-C. **(E)** Scatter plots showing the correlation between CHiCAGO (inverse hyperbolic sine-transformed) and Peaky (square root-transformed) interaction scores. ? is Spearman’s correlation; the black lines represent the loess smoothed fit. **(F)** Venn diagrams illustrating the overlap in CHiCAGO-and Peaky-scored interactions in each capture per cell line.

**Figure S4.**
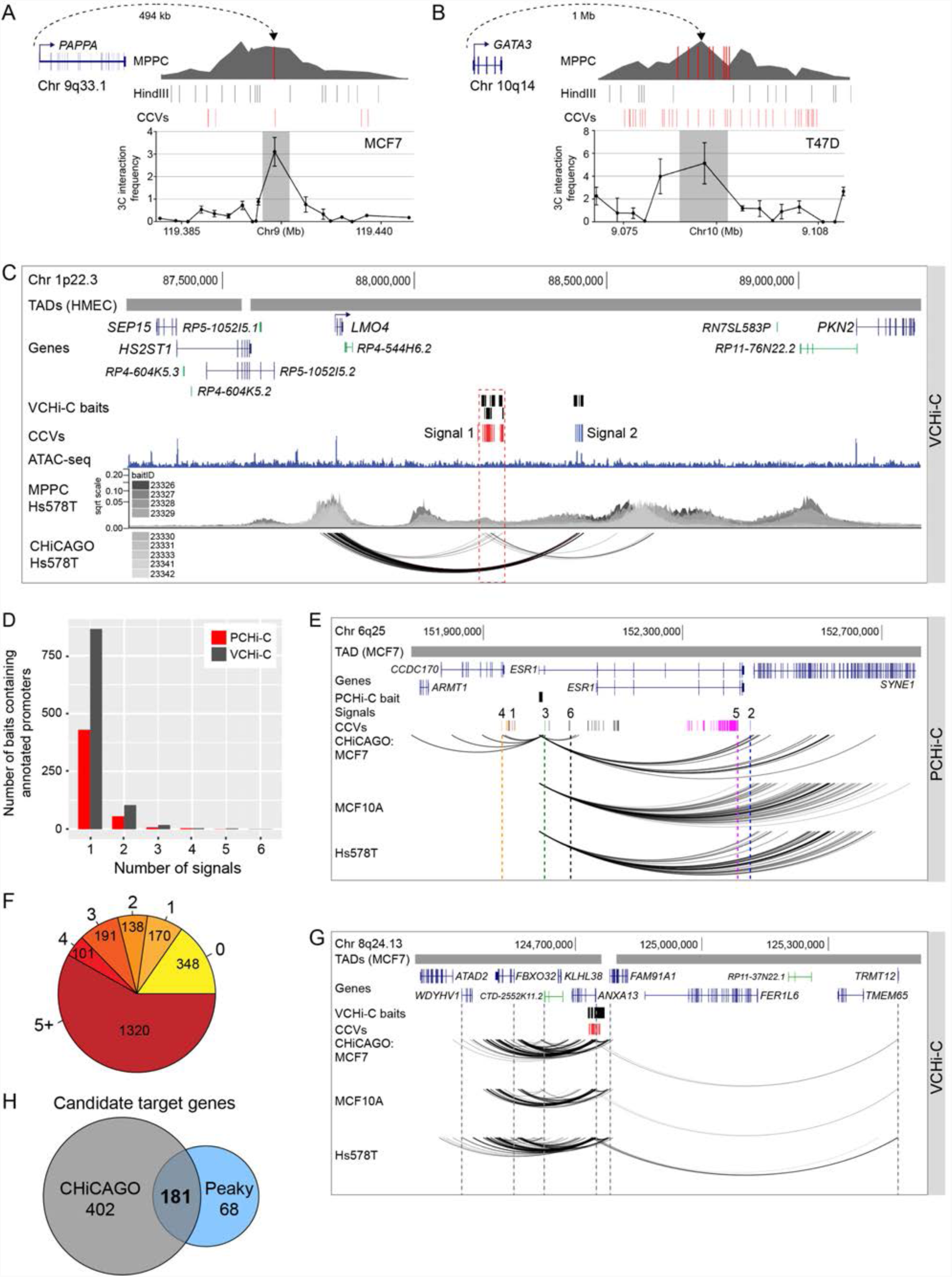
Candidate target gene properties, related to Figures 3 and 4. **(A)** 3C interaction profiles at 9q33.1 between the *PAPPA* promoter and CCVs in MCF7 cells. The anchor point is set at the *PAPPA* promoter. Error bars represent SD (n=3). **(B)** 3C interaction profiles at 10q14 between the *GATA3* promoter and CCVs in T47D cells. The anchor point is set at the *GATA3* promoter. Error bars represent SD (n=3). **(C)** Chromatin interactions at 1p22.3 in Hs578T cells. Topologically associating domains (TADs) are shown as horizontal gray bars above GENCODE annotated coding (blue) and non-coding (green) genes. The VCHi-C baits are depicted as black boxes. Risk signals 1 and 2 are numbered and the CCVs within each signal are shown as colored vertical lines. The ATAC-seq track is shown as a dark blue histogram. Peaky defined MPPC values (from nine specified BaitIDs) are plotted. CHiCAGO-scored interactions are shown as black arcs. The dashed red outline highlights the signal 1 CCVs. **(D)** Correlation between the number of candidate target genes and independent risk signals in the PCHi-C and VCHi-C datasets. **(E)** Chromatin interactions at 6q25 in MCF7, MCF10A and Hs578T breast cell lines. Topologically associating domains (TADs) are shown as horizontal gray bars above GENCODE annotated coding (blue) genes. The *ESR1* PCHi-C bait (BaitID: 355261) is depicted as a black box. Risk signals 1-6 are numbered and the CCVs within each signal are shown as colored vertical lines. CHiCAGO-scored interactions are shown as black arcs. The dashed colored vertical lines highlight *ESR1* promoter-signal interactions. **(F)** Enumeration of the number of transcription start sites skipped during chromatin looping for the 651 target gene promoter interactions. **(G)** Chromatin interactions at 8q24.13 in MCF7, MCF10A and Hs578T breast cell lines. Topologically associating domains (TADs) are shown as horizontal gray bars above GENCODE annotated coding (blue) and non-coding (green) genes. VCHi-C baits are depicted as black boxes and the signal 1 CCVs as red vertical lines. CHiCAGO-scored interactions are shown as black arcs. The dashed gray vertical lines highlight promoter-signal interactions to the candidate target genes. **(H)** Venn diagram illustrating the number of candidate target genes identified by CHiCAGO, Peaky and both algorithms.

**Figure S5.**
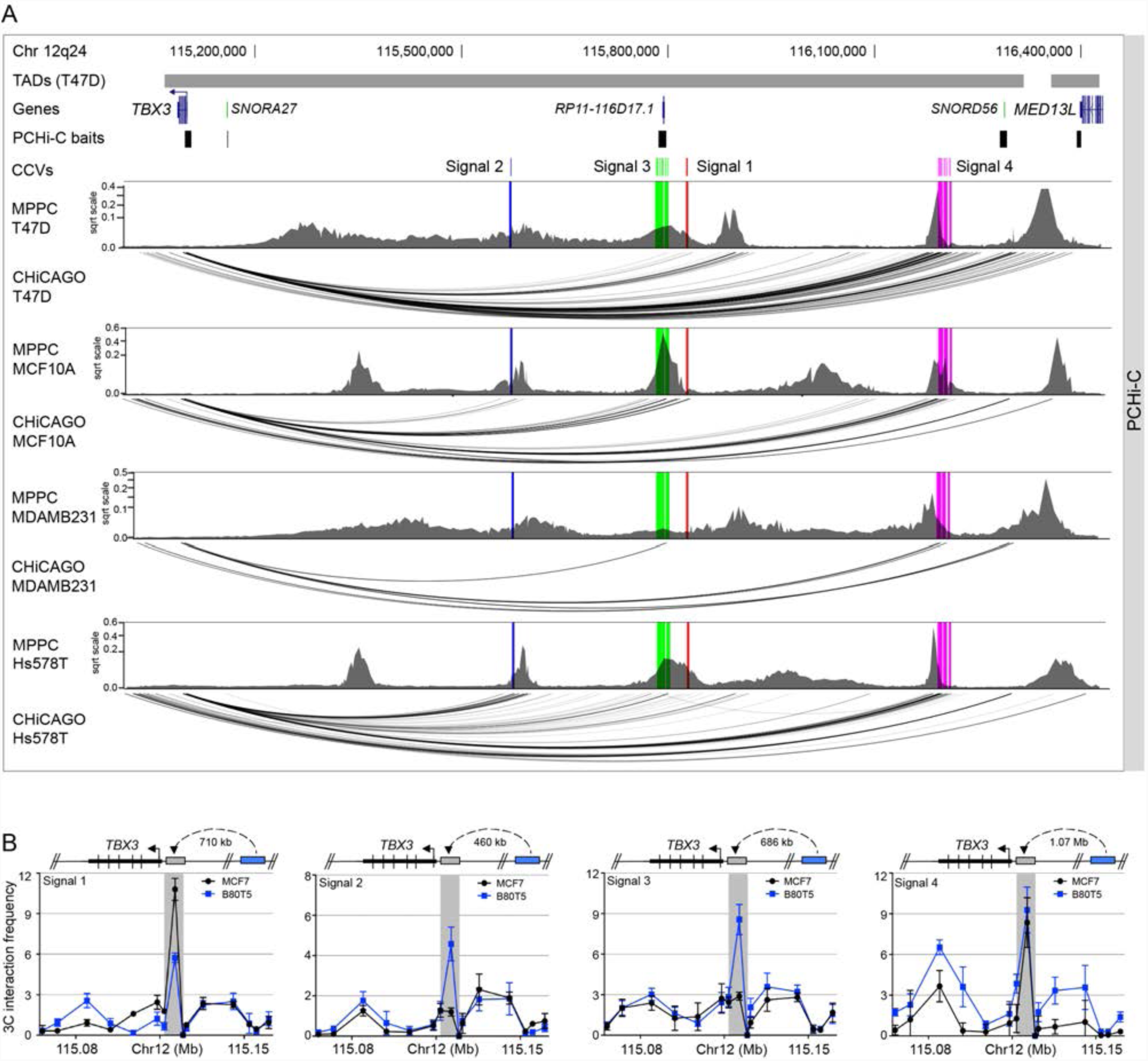
Chromatin interactions across 12q24, related to Figure 5. **(A)** Chromatin interactions at 12q24 in ER+ T47D, ER-MDAMB231 and Hs578T breast cancer cell lines, and MCF10A non-tumorigenic breast cells. Topologically associating domains (TADs) are shown as horizontal gray bars above GENCODE annotated coding (blue) and non-coding (green) genes. PCHi-C baits are depicted as black boxes. Risk signals 1-4 are numbered and the CCVs within each signal are shown as colored vertical lines. Peaky defined MPPC values (from PCHi-C baitID 596031) are plotted with the CCVs overlaid as colored vertical lines. CHiCAGO-scored interactions are shown as black arcs. **(B)** 3C interaction profiles between the risk signals 1-4 and the *TBX3* promoter in MCF7 and B80T5 cell lines. Anchor points are set at signals 1-4. Error bars represent SD (n=3).

**Figure S6.**
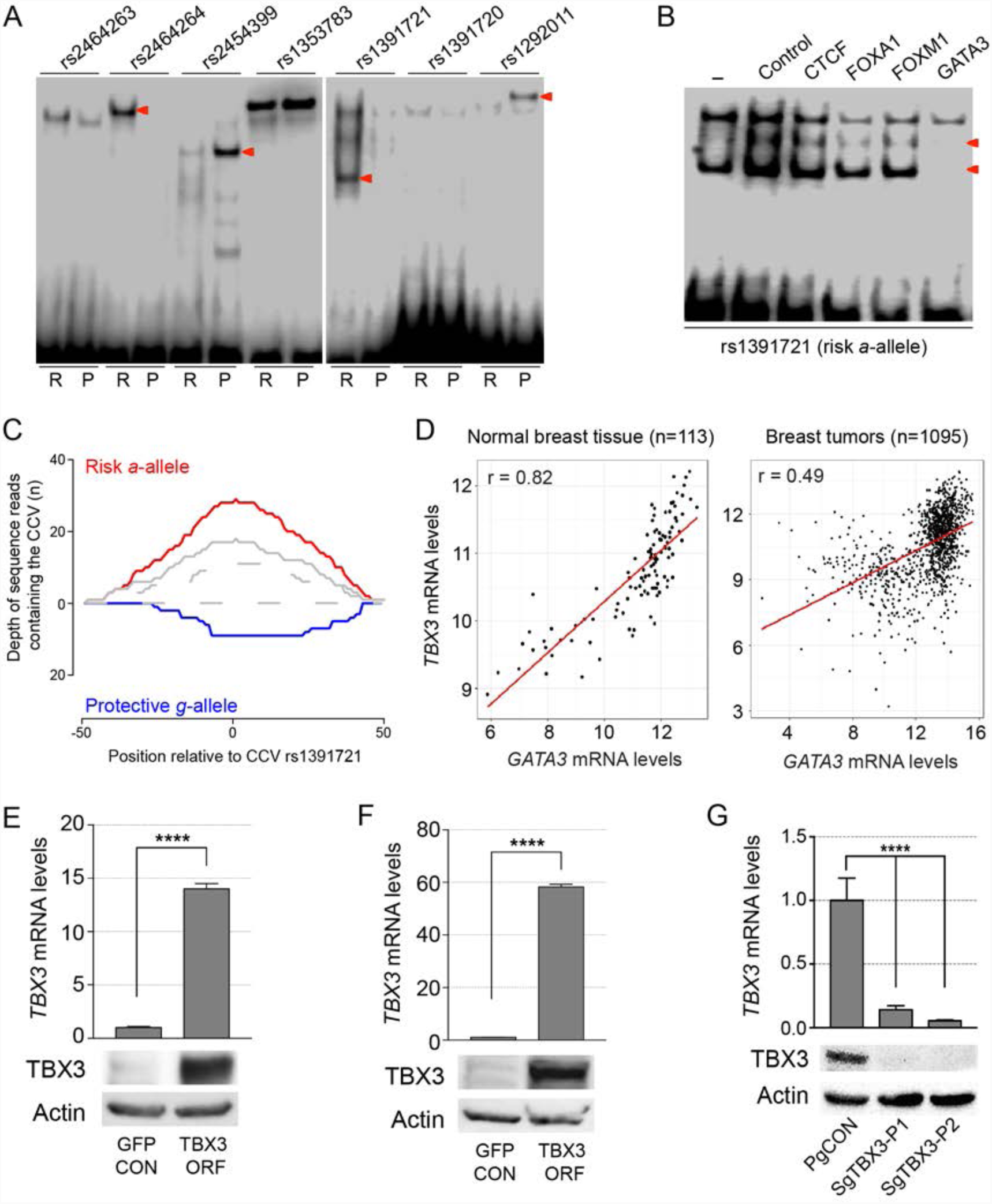
Additional *in vitro* and *in vivo* studies for 12q24, related to Figures 5 and 6. **(A)** EMSAs for signal 1 CCVs to detect allele-specific binding of nuclear proteins. Labeled oligonucleotide duplexes were incubated with BT474 nuclear extract. Red arrowheads show bands of different mobility detected between risk (R) and protective (P) alleles. **(B)** EMSAs for CCV rs1391721 to identify candidate nuclear proteins. Unlabeled competitor oligonucleotide duplexes for predicted transcription factors (100-fold molar excess) were incubated with labeled rs1391721-containing oligonucleotide duplex and MCF7 nuclear extract. Red arrowheads indicate bands that were competed for complex formation on the risk (*a*) allele. **(C)** Allele-specific GATA3 binding at CCV rs1391721 in heterozygous MCF7 cells. The depth of reads containing the risk (red) and protective (blue) alleles are shown. **(D)** Scatter plots of *TBX3* versus *GATA3* gene expression in TCGA normal breast tissue (n=113, r is Pearson’s correlation) and breast tumors (n=1095, r is Pearson’s correlation). **(E)** Top: *TBX3* levels in HMLE-control (GFP CON) and HMLE-TBX3 overexpressing (TBX3 ORF) cells assessed by qPCR and normalized to *GUSB*. Error bars represent SEM (n=3). Bottom: Western blot analysis of TBX3 and Actin, serving as a loading control, in matched cell samples. **(F)** Top: *TBX3* levels in MCF7-control (GFP CON) and MCF7-TBX3 overexpressing (TBX3 ORF) cells assessed by qPCR and normalized to *GUSB*. Error bars represent SEM (n=3). Bottom: Western blot analysis of TBX3 and Actin in matched cell samples. **(G)** Top: TBX3 levels in MCF7-control (PgCON) and MCF7-TBX3-dCas9-KRAB repressed cells (SgTBX3-P1/P2) assessed by qPCR and normalized to *GUSB*. Error bars represent SEM (n=3). Bottom: Western blot analysis of TBX3 and Actin in matched cell samples. **(E-G)** *P*-values were determined by a two-tailed *t*-test (****p<0.0001).

## REFERENCES

1. Melchor, L. & Benitez, J. The complex genetic landscape of familial breast cancer. Hum Genet 132, 845–63 (2013).

2. Michailidou, K. et al. Association analysis identifies 65 new breast cancer risk loci. Nature 551, 92–94 (2017).

3. Mavaddat, N. et al. Prediction of breast cancer risk based on profiling with common genetic variants. J Natl Cancer Inst 107(2015).

4. Edwards, S.L., Beesley, J., French, J.D. & Dunning, A.M. Beyond GWASs: illuminating the dark road from association to function. Am J Hum Genet 93, 779–97 (2013).

5. Fachal, L. et al. Fine mapping of 150 breast cancer risk regions identifies 178 high confidence target genes. Submitted.

6. Cavalli, G. & Misteli, T. Functional implications of genome topology. Nat Struct Mol Biol 20, 290– 9 (2013).

7. Bonev, B. & Cavalli, G. Organization and function of the 3D genome. Nat Rev Genet 17, 772 (2016).

8. Nora, E.P. et al. Spatial partitioning of the regulatory landscape of the X-inactivation centre. Nature 485, 381–5 (2012).

9. Dixon, J.R. et al. Topological domains in mammalian genomes identified by analysis of chromatin interactions. Nature 485, 376–80 (2012).

10. Naumova, N., Smith, E.M., Zhan, Y. & Dekker, J. Analysis of long-range chromatin interactions using Chromosome Conformation Capture. Methods 58, 192–203 (2012).

11. Lieberman-Aiden, E. et al. Comprehensive mapping of long-range interactions reveals folding principles of the human genome. Science 326, 289–93 (2009).

12. Rao, S.S. et al. A 3D map of the human genome at kilobase resolution reveals principles of chromatin looping. Cell 159, 1665–80 (2014).

13. Jin, F. et al. A high-resolution map of the three-dimensional chromatin interactome in human cells. Nature 503, 290–4 (2013).

14. Dryden, N.H. et al. Unbiased analysis of potential targets of breast cancer susceptibility loci by Capture Hi-C. Genome Res 24, 1854–68 (2014).

15. Schoenfelder, S. et al. The pluripotent regulatory circuitry connecting promoters to their long-range interacting elements. Genome Res 25, 582–97 (2015).

16. Javierre, B.M. et al. Lineage-specific genome architecture links enhancers and non-coding disease variants to target gene promoters. Cell 167, 1369–1384 e19 (2016).

17. Rubin, A.J. et al. Lineage-specific dynamic and pre-established enhancer-promoter contacts cooperate in terminal differentiation. Nat Genet 49, 1522–1528 (2017).

18. Mifsud, B. et al. Mapping long-range promoter contacts in human cells with high-resolution capture Hi-C. Nat Genet 47, 598–606 (2015).

19. Davies, J.O. et al. Multiplexed analysis of chromosome conformation at vastly improved sensitivity. Nat Methods 13, 74–80 (2016).

20. Siersbaek, R. et al. Dynamic rewiring of promoter-anchored chromatin loops during adipocyte differentiation. Mol Cell 66, 420–435 e5 (2017).

21. McGovern, A. et al. Capture Hi-C identifies a novel causal gene, IL20RA, in the pan-autoimmune genetic susceptibility region 6q23. Genome Biol 17, 212 (2016).

22. Martin, P. et al. Capture Hi-C reveals novel candidate genes and complex long-range interactions with related autoimmune risk loci. Nat Commun 6, 10069 (2015).

23. Cairns, J. et al. CHiCAGO: robust detection of DNA looping interactions in Capture Hi-C data. Genome Biol 17, 127 (2016).

24. Roadmap Epigenomics, C. et al. Integrative analysis of 111 reference human epigenomes. Nature 518, 317–30 (2015).

25. Theodorou, V., Stark, R., Menon, S. & Carroll, J.S. GATA3 acts upstream of FOXA1 in mediating ESR1 binding by shaping enhancer accessibility. Genome Res 23, 12–22 (2013).

26. Stevens, T.J. et al. 3D structures of individual mammalian genomes studied by single-cell Hi-C. Nature 544, 59–64 (2017).

27. Curtis, C. et al. The genomic and transcriptomic architecture of 2,000 breast tumours reveals novel subgroups. Nature 486, 346–52 (2012).

28. Williamson, I. et al. Spatial genome organization: contrasting views from chromosome conformation capture and fluorescence in situ hybridization. Genes Dev 28, 2778–91 (2014).

29. Rosa, A., Becker, N.B. & Everaers, R. Looping probabilities in model interphase chromosomes. Biophys J 98, 2410–9 (2010).

30. Eijsbouts, C., Burren, O.S., Newcombe, P. & Wallace, C. Fine mapping chromatin contacts in capture Hi-C data. bioRxiv (2018).

31. Conover, C.A. Key questions and answers about pregnancy-associated plasma protein-A. Trends Endocrinol Metab 23, 242–9 (2012).

32. Mansfield, A.S. et al. Pregnancy-associated plasma protein-A expression in human breast cancer. Growth Horm IGF Res 24, 264–7 (2014).

33. Takabatake, Y. et al. Lactation opposes pappalysin-1-driven pregnancy-associated breast cancer. EMBO Mol Med 8, 388–406 (2016).

34. Visvader, J.E. et al. The LIM domain gene LMO4 inhibits differentiation of mammary epithelial cells in vitro and is overexpressed in breast cancer. Proc Natl Acad Sci U S A 98, 14452–7 (2001).

35. Sum, E.Y. et al. Overexpression of LMO4 induces mammary hyperplasia, promotes cell invasion, and is a predictor of poor outcome in breast cancer. Proc Natl Acad Sci U S A 102, 7659–64 (2005).

36. Hannenhalli, S. & Kaestner, K.H. The evolution of Fox genes and their role in development and disease. Nat Rev Genet 10, 233–40 (2009).

37. Mani, S.A. et al. Mesenchyme Forkhead 1 (FOXC2) plays a key role in metastasis and is associated with aggressive basal-like breast cancers. Proc Natl Acad Sci U S A 104, 10069–74 (2007).

38. Zhong, J., Wang, H., Yu, J., Zhang, J. & Wang, H. Overexpression of Forkhead Box L1 (FOXL1) inhibits the proliferation and invasion of breast cancer cells. Oncol Res 25, 959–965 (2017).

39. MacNair, L. et al. MTHFSD and DDX58 are novel RNA-binding proteins abnormally regulated in amyotrophic lateral sclerosis. Brain 139, 86–100 (2016).

40. Nik-Zainal, S. et al. Landscape of somatic mutations in 560 breast cancer whole-genome sequences. Nature 534, 47–54 (2016).

41. ENCODE Project Consortium. An integrated encyclopedia of DNA elements in the human genome. Nature 489, 57–74 (2012).

42. Willmer, T., Cooper, A., Peres, J., Omar, R. & Prince, S. The T-Box transcription factor 3 in development and cancer. Biosci Trends 11, 254–266 (2017).

43. Elenbaas, B. et al. Human breast cancer cells generated by oncogenic transformation of primary mammary epithelial cells. Genes Dev 15, 50–65 (2001).

44. Willmer, T., Cooper, A., Sims, D., Govender, D. & Prince, S. The T-box transcription factor 3 is a promising biomarker and a key regulator of the oncogenic phenotype of a diverse range of sarcoma subtypes. Oncogenesis 5, e199 (2016).

45. Dunning, A.M. et al. Breast cancer risk variants at 6q25 display different phenotype associations and regulate ESR1, RMND1 and CCDC170. Nat Genet 48, 374–86 (2016).

46. Meyer, K.B. et al. Fine-scale mapping of the FGFR2 breast cancer risk locus: putative functional variants differentially bind FOXA1 and E2F1. Am J Hum Genet 93, 1046–60 (2013).

47. Ghoussaini, M. et al. Evidence that breast cancer risk at the 2q35 locus is mediated through IGFBP5 regulation. Nat Commun 4, 4999 (2014).

48. Baxter, J.S. et al. Capture Hi-C identifies putative target genes at 33 breast cancer risk loci. Nat Commun 9, 1028 (2018).

49. Maurano, M.T. et al. Systematic localization of common disease-associated variation in regulatory DNA. Science 337, 1190–5 (2012).

50. Fischer, K. & Pflugfelder, G.O. Putative Breast Cancer Driver Mutations in TBX3 Cause Impaired Transcriptional Repression. Front Oncol 5, 244 (2015).

51. Shen, L., Shi, Q. & Wang, W. Double agents: genes with both oncogenic and tumor-suppressor functions. Oncogenesis 7, 25 (2018).

52. Nagano, T. et al. Comparison of Hi-C results using in-solution versus in-nucleus ligation. Genome Biol 16, 175 (2015).

53. Wingett, S. et al. HiCUP: pipeline for mapping and processing Hi-C data. F1000Res 4, 1310 (2015).

54. Buenrostro, J.D., Giresi, P.G., Zaba, L.C., Chang, H.Y. & Greenleaf, W.J. Transposition of native chromatin for fast and sensitive epigenomic profiling of open chromatin, DNA-binding proteins and nucleosome position. Nat Methods 10, 1213–8 (2013).

55. Martin, M. Cutadapt removes adapter sequences from high-throughput sequencing reads. 2011 17(2011).

56. Li, H. & Durbin, R. Fast and accurate long-read alignment with Burrows-Wheeler transform. Bioinformatics 26, 589–95 (2010).

57. Li, H. et al. The Sequence Alignment/Map format and SAMtools. Bioinformatics 25, 2078–9 (2009).

58. Zhang, Y. et al. Model-based analysis of ChIP-Seq (MACS). Genome Biol 9, R137 (2008).

59. Heinz, S. et al. Simple combinations of lineage-determining transcription factors prime cis-regulatory elements required for macrophage and B cell identities. Mol Cell 38, 576–89 (2010).

